# Genomes of the human filarial parasites *Mansonella perstans* and *Mansonella ozzardi*

**DOI:** 10.1101/2023.01.06.523030

**Authors:** Amit Sinha, Zhiru Li, Catherine B. Poole, Richard D. Morgan, Laurence Ettwiller, Nathália F. Lima, Marcelo U. Ferreira, Fanny F. Fombad, Samuel Wanji, Clotilde K. S. Carlow

## Abstract

The filarial parasites *Mansonella ozzardi* and *Mansonella perstans*, causative agents of mansonellosis, infect hundreds of millions of people worldwide, yet remain among the most understudied of the human filarial pathogens. *M. ozzardi* is highly prevalent in Latin American countries and Caribbean Islands, while *M. perstans* is predominantly found in sub-Saharan Africa as well as in a few areas in South America. In addition to the differences in their geographical distribution, the two parasites are transmitted by different insect vectors, as well as exhibit differences in their responses to commonly used anthelminthic drugs. The lack of genome information has hindered investigations into the biology and evolution of *Mansonella* parasites and understanding the molecular basis of the clinical differences between species. In the current study, high quality genomes of two independent clinical isolates of *M. perstans* from Cameroon and two *M. ozzardi* isolates one from Brazil and one from Venezuela are reported. The genomes are approximately 76 Mb in size, encode about 10,000 genes each, and are largely complete based on BUSCO scores of about 90%, similar to other completed filarial genomes. These sequences represent the first genomes from *Mansonella* parasites and enabled a comparative genomic analysis of the similarities and differences between *Mansonella* and other filarial parasites. Horizontal DNA transfers (HDT) from mitochondria (nuMTs) as well as transfers from genomes of endosymbiotic *Wolbachia* bacteria (nuWTs) to the host nuclear genome were identified and analyzed. Sequence comparisons and phylogenetic analysis of known targets of anti-filarial drugs diethylcarbamazine (DEC), ivermectin and mebendazole revealed that all known target genes were present in both species, except for the DEC target encoded by *gon-2* gene, which is fragmented in genome assemblies from both *M. ozzardi* isolates. These new reference genome sequences will provide a valuable resource for further studies on biology, symbiosis, evolution and drug discovery.

## 1 Introduction

A large part of the world’s population is infected with one or more filarial nematode parasites resulting in several distinct diseases often referred to as a group under the general term filariasis. The filarial parasites *Mansonella ozzardi* and *Mansonella perstans* are the two main causes of mansonellosis in humans. Infections are transmitted by infected biting midges (mostly of the genus *Culicoides*) or blackflies (genus *Simulium*) that release infective stage larvae during a blood meal. Adult male and female worms have been observed rarely in humans since they are presumed to live in the serous body cavities and/or subcutaneous tissues. Viviparous females release unsheathed microfilaria which circulate in the blood, available for ingestion by an appropriate vector during feeding. Unlike some of the other human filarial diseases, *Mansonella* infections do not present with a distinct pathology or specific clinical picture. Infected individuals are often asymptomatic or may have a variety of clinical features including subcutaneous swellings, aches, pain, fever, headache, pruritis, corneal lesions and eosinophilia (Simonsen et al., 2011; Lima et al., 2016; Mediannikov & Ranque 2018; Vianna et al., 2012; Ferreira et al 2021). Other manifestations include immunosuppression (Ritter et al., 2018) with important ramifications such as increased susceptibility to other infections and reduced efficacy of vaccine programs (Ta-Tang et al., 2018).

Even though mansonellosis ranks first in prevalence among the human filariases, affecting hundreds of millions of people in Africa and Central and South America, it remains one of the least studied and most neglected of all filarial diseases (Downes & Jacobsen, 2010; Simonsen et al., 2011; Lima et al., 2016; Mediannikov & Ranque, 2018; Ta-Tang et al., 2018; Ta-Tang et al., 2021; Sandri et al., 2021; Bélard & Gehringer 2021). Indeed, much of the somewhat limited information available has been obtained serendipitously during studies on other filarial parasites or malaria, in which individuals were co-infected with *Mansonella* parasites. *M. perstans* is considered the most common filariasis in Central Africa with infection prevalence often very high (up to 96%) in endemic areas, even among children (Downes and Jacobsen, 2010; Simonsen et al., 2011; TaTang et al., 2018). It is also found in northern regions of the Amazon rainforest and the Caribbean coast in South America (TaTang et al., 2018; Bélard and Gehringer, 2021), with molecular evidence supporting introduction of *M. perstans* to the American continent from Africa because of the slave trade (Tavares da Silva et al., 2017). *M. ozzardi* infections have been reported from many countries in South America, Central America, and some Caribbean Islands (Lima et al., 2018; TaTang et al.,2018; Tavares de Silva et al., 2017; Raccurt et al., 2018; Ferriera et al., 2021). Co-infections of *M. perstans* and *M. ozzardi*, together with other filarial parasites, have also been reported in South America (Kozek et al., 1983; Calvopina et al., 2019), making their treatment challenging as the two species respond differently to the commonly used anti-filarial drugs such as ivermectin, diethylcarbamazine (DEC) or mebendazole (Chadee et al., 1995; TaTang et al., 2018). The recent development of highly specific, simple molecular tests which can distinguish *M. perstans* and *M. ozzardi*, and do not cross-react with other filarial parasites (Poole et al., 2019), will facilitate an effective treatment regimen. The antibiotic doxycycline, targeting the endosymbiotic *Wolbachia* bacteria of *M. perstans* and *M. ozzardi*, has been shown to clear the microfilarial stage from the blood (Coulibaly et al., 2009; Debrah et al., 2019; Crainey et al., 2020) highlighting the importance of *Wolbachia* in these species. The *Wolbachia* genomes, *w*Mpe from *M. perstans*, and *w*Moz from *M. ozzardi* have recently been published (Sinha et al., 2022), providing a valuable resource for the discovery of new antibiotics and studies on the symbiotic relationship and evolution.

Here we provide the first *de novo* assemblies of the nuclear genomes from *M. perstans* and *M. ozzardi* using combined PacBio long-read and Illumina paired-end sequencing data. We used customized metagenomic assembly and binning pipelines, and extensive quality checks to ensure accurate assemblies from a mixture of DNA from the *Mansonella* parasites, their human host, and the associated microbiome. These methods were applied to generate genomes from two clinical isolates of each species, providing a unique opportunity for intra-species and inter-species comparisons and exploring features of sequence evolution of these parasites. We analyzed the genomic regions containing horizontally transferred DNA from mitochondria (nuMTs), as well horizontally transferred DNA from their endosymbiont *Wolbachia* (nuWTs). Phylogenetic analysis of orthologs of genes that encode drug targets of common anti-filarial drugs such as DEC, ivermectin and mebendazole revealed intriguing differences in the presence and absence of these targets between *M. perstans* and *M. ozzardi* that might have clinical implications for the treatment of mansonellosis.

## 2 Methods

### 2.1 Ethics statement

All research involving human subjects was approved by the appropriate committee and performed in accordance with all relevant guidelines and regulations. Informed consent was obtained from all participants or their legal guardians. These samples have also been described and utilized in previous studies (Poole et al., 2019; Sinha et al., 2022).

Ethical clearance for *M. perstans* samples was obtained from the National Institutional Review board, Yaoundé (REF: N° 2015/09/639/CE/CNERSH/SP) and administrative clearance from the Delegation of Public Health, South-West region of Cameroon (Re: R11/MINSANTE/SWR/ RDPH/PS/259/382). Approval for the study was granted by the “National Ethics Committee of Research for Human Health” in Cameroon. Special consideration was taken to minimize the health risks to which any participant in this study was exposed. The objectives of the study were explained to the consenting donor after which they signed an informed consent form. The participant’s documents were given a code to protect the privacy of the study subject. At the end of the study, the donor received a cure of mebendazole (100 mg twice a day for 30 days).

The study protocols for *M. ozzardi* were approved by the Institutional Review Board of the Institute of Biomedical Sciences, University of São Paulo, Brazil (1133/CEP, 2013) as part of a previous study (Lima et al., 2018). Written informed consent was obtained from all patients, or their parents or guardians if participants were minors younger than 18 years. Diagnosed infections were treated with a single dose of 0.2 mg/kg of ivermectin after blood sampling (Lima et al., 2018).

### 2.2 Illumina library construction and sequencing

The parasite materials, DNA extractions, Illumina library preparation and sequencing have been described in detail previously (Poole et al., 2019; Sinha et al., 2022). Briefly, two clinical isolates of *M. perstans*, named Mpe-Cam-1 and Mpe-Cam-2, were obtained as microfilariae in a blood draw from consenting individuals in Cameroon. One clinical isolate of *M. ozzardi*, named Moz-Brazil-1, was obtained from a consenting individual as part of a previous study in Brazil, while another isolate from Venezuela, named Moz-Venz-1, was generously donated by Izaskun Petralanda in 1989. The DNA samples from Mpe-Cam-1, Mpe-Cam-2, and Moz-Brazil-1 were treated with the NEBNext Microbiome DNA enrichment to reduce human DNA contamination before being used for library preparation. Illumina libraries for all 4 isolates were constructed using the NEBNext Ultra II FS DNA Library Prep Kit (New England Biolabs Inc., USA) as per standard protocol and sequenced using the Illumina MiSeq and NextSeq500 platforms (paired end, 150 bps). Bioinformatic analysis programs were run with their default parameters unless otherwise stated. The bioinformatic analysis pipeline and scripts are available from https://github.com/aWormGuy/Mansonella-Genomes-Sinha-et-al.-2023.

### 2.3 PacBio library sequencing and *de novo* assembly

A total of 250 ng of genomic DNA from *M. perstans* isolate Mpe-Cam-1 was used for sequencing on PacBio RSII platform as per manufacturer’s instructions. Consensus reads from raw PacBio reads were obtained using the protocol RS_PreAssembler.1 (parameters minLen=1000, minQual=0.80, genomeSize=100000000) on PacBio SMRT Portal (smrtanalysis_2.3.0.140936.p5.167094). Reads originating from human genomic DNA were removed by first by mapping the consensus reads to the human genome (grch38) using minimap2 v2.17-r941 (Li, 2018) followed by discarding all mapped reads (MAPQ score > 20). The remaining reads were assembled using canu v2.2 (Koren et al., 2017) (parameters genomeSize=90m correctedErrorRate=0.045). The assembly was iteratively polished over 12 rounds using “Resequencing” protocol on PacBio SMRT Portal (v 6.0.0.47841) (parameters --minMatch 12 --bestn 10 --minPctSimilarity 70.0 --refineConcordantAlignments) till no new variants could be detected by the Resequencing pipeline. The assembly contiguity was further improved by using finisherSC software v2.1 (Lam et al., 2015), a scaffolder that uses raw PacBio reads while accounting for any genomic repeats in the assembly. The assembly was polished using the Illumina data for the same isolate via the polca software (Zimin and Salzberg, 2020). Finally, gap-filling and heterozygosity removal was performed on this assembly using Illumina reads as an input to the Redundans pipeline v0.14a (Pryszcz and Gabaldón, 2016). The assembly quality was evaluated using quast (Gurevich et al., 2013) at each step.

### 2.4 Metagenomic assembly using Illumina reads

Illumina raw reads were processed using the BBTools package (https://jgi.doe.gov/data-and-tools/bbtools/). Duplicate raw reads and bad tiles were removed using the clumpify.sh and filterbytile.sh utilities. Adapter trimming, removal of phiX reads, and reads with 13-nt or longer stretches of the same nucleotide (poly-G, poly-C, poly-A, poly-T) was performed using bbduk.sh. Human host reads were removed by mapping against the human genome (grch38) using bbmap.sh utility. The quality metrics of the processed reads at each step were assessed using FastqC (https://www.bioinformatics.babraham.ac.uk/projects/fastqc/). Reads from different runs of the libraries prepared from the same genomic DNA sample were combined and used as an input for the assembly of the metagenome using metaSpades (Nurk et al., 2017).

### 2.5 Metagenomic binning to identify *Mansonella* contigs

Binning of metagenomic contigs to identify sequences originating from *Mansonella* genomes was carried out based on the BlobTools software (Laetsch and Blaxter, 2017) and additional curation, as described previously (Sinha et al., 2022). Briefly, for each of the isolates Mpe-Cam-2, Moz-Brazil-1 and Moz-Venz-1, the Illumina reads were mapped back to the metagenomic contigs produced by metaSpades using bowtie2 (Langmead and Salzberg 2012). For the PacBio assembly obtained from the Mpe-Cam-1 sample, the corresponding Illumina reads were mapped to this assembly using bowtie2. Sequence similarities of assembled contigs to the NCBI nt database and Uniport database was carried out using blastn and diamond in BlobTools package. The bins annotated as “Nematoda” and “Proteobacteria” by BlobTools were analyzed further to retrieve sequences originating from *Mansonella*. Contigs that were marked as Proteobacteria but co-localized with the Nematoda blobs in the blobplots, were included in the *Mansonella* bins as they represented potential examples of horizontal gene transfer from *Wolbachia* to *Mansonella* host. These HGT candidates were validated by inspection of blastn and blastx hits of these contigs to the NCBI nt and nr databases respectively.

Contigs originating from mitochondrial DNA were identified by analysis of nucmer alignments between assembled sequences and the previously reported mitochondrial genomes, GenBank accessions MT361687.1 and KX822021.1 for *M. perstans* (Chung et al., 2020) and *M. ozzardi* (Crainey et al., 2018) respectively. The contigs with more than 98% sequence identity to published mitochondrial genomes were marked as mtDNA and removed from the nuclear assembly.

### 2.6 Assembly refinements and heterozygosity removal for Illumina assemblies

For each Illumina-only assembly, scaffolding, gap-filling and heterozygosity removal was performed using the corresponding Illumina reads in the Redundans pipeline v0.14a (Pryszcz and Gabaldón, 2016).

### 2.7 Genome annotations and comparative analysis

Identification of repetitive elements in the assembled genomes were performed using RepeatModeler version 2.0.3 (Flynn et al., 2020) and classified against the Dfam database (Storer et al., 2021). First round of repeat annotations was performed on a fasta file combining all assembled genomes. The repeat sequence families identified from this run were used as a reference library in RepeatMasker 4.1.2-p1 (http://repeatmasker.org/) to mask repeats in each of the 4 genome assemblies independently. The outputs from RepeatMasker were processed with parseRM.pl utility (Kapusta et al. 2017) to generate summary statistics on repats classified as DNA transposons and retroelements.

Gene predictions were carried out using the braker2 pipeline v2.2.6 (Bruna et al., 2020), with a reference proteome comprised of all metazoan proteins in OrthoDB version 10.1 (Kriventseva et al., 2019) and all filarial proteomes downloaded from WormBase ParaSite (Howe et al., 2017). To ensure that the same parameters and hmm models are applied to all assemblies, the 4 assemblies were combined into a single file, and braker2 pipeline was run on this composite dataset. The genes predicted from this run were then assigned to respective isolates by parsing the gff3 output from braker2. The completeness of the protein-coding content of genomes was assessed using the BUSCO pipeline v5.0 beta using the “nematoda_odb10” reference dataset (Simão et al., 2015). OrthoFinder v2.4 (Emms and Kelly, 2019) was used to infer orthologs of protein coding genes across multiple nematode genomes. The set of proteomes included all 17 filarial species with publicly available genomes. The species *Setaria digitata* and *Thelazia callipaeda* from the same infraorder Spiruromorpha were included as the immediate outgroups, while *Caenorhabditis elegans* and *Pristionchus pacificus* were included as relatively more distant outgroups. The protein and transcript sequences were downloaded from WormBase parasite where available. Although the genome sequences for filarial nematodes *Cruorifilaria tuberocauda*, *Dipetalonema caudispina*, *Dirofilaria immitis*, *Litomosoides brasiliensis* and *Madathamugadia hiepei* (Lefoulon et al., 2020) were available from NCBI (GenBank accessions GCA_013365365.1, GCA_013365325.1, GCA_013365355.1, GCA_013365375.1 and GCA_013365335.1 respectively), the corresponding gene annotations for these assemblies were not available in NCBI. Hence, repeat-modeling, repeat-masking and gene annotation using braker2 pipeline was performed on each of these species as described for *M. perstans* and *M. ozzardi* above, and have been made available at https://github.com/aWormGuy/Mansonella-Genomes-Sinha-et-al.-2023/tree/main/gene_predictions_on_Lefoulon2020_genomes. available a. The source databases, the corresponding accession numbers, filenames, citations and BUSCO scores for all the genomes used in comparative analysis are listed in Supplementary Material Table 1.

Whole-genome alignments for global sequence similarity and synteny between various repeat-masked genomes were performed using minimap2 v2.17-r941 (Li, 2018) and visualized using the JupiterPlot tool (https://github.com/JustinChu/JupiterPlot). For global sequence comparisons across multiple filarial genomes, the average nucleotide identity (ANI) scores were calculated using the OrthoANIu tool (Yoon et al., 2017).

Comparison and analysis of sequence differences, including structural difference, between different pairs of genomes, namely (A) Mpe-Cam-1versus Mpe-Cam-2 (B) Moz-Brazil-1 versus Moz-Venz-1 (C) Mpe-Cam-1 versus Moz-Brazil-1 were performed using nucdiff version 2.0.3 (Khelik et al., 2017). The localization of the sequence variants within gene bodies and exons was calculated through the intersectBed tool in the bedtools package v2.29.2 (Quinlan and Hall, 2010). The effect of sequence variants on the coding potential of affected genes was determined using snpEff version 5.1d (Cingolani et al., 2012).

### 2.8 Phylogenetic analysis

For genome-wide phylogenetic analysis, single copy orthologs across various nematodes were identified from the OrthoFinder analysis described above. The corresponding protein sequences were aligned using mafft v7.149b (Nakamura et al., 2018) and trimmed using Gblocks v0.91b (Talavera and Castresana, 2007). Phylogenetic analysis of the resulting supermatrix of 823,832 amino acids was carried out using iqTree v2.1.2 (Nguyen et al., 2015). The best fit substitution model was chosen automatically using ModelFinder implementation within iqTree (Kalyaanamoorthy et al., 2017). Bootstrap support values were calculated based on (i) the SH-like approximate likelihood ratio test (Guindon et al., 2010), and (ii) ultrafast bootstrap (Hoang et al., 2018) with 1000 replicates each (iqTree command-line options ‘-B 1000 -alrt 1000 -abayes -lbp 1000’).

For phylogenetic analysis of specific genes and gene families, their homologs and the corresponding protein sequences were obtained from the OrthoFinder output. The protein domains for all the sequences were annotated using NCBI CD search and the NCBI CDD database (Lu et al., 2020). The protein sequences which lacked the domains present in the *C. elegans* homologs, or were too short in length (less than 60% of the length of the *C. elegans* homolog) were inspected and excluded from the final analysis. The multiple sequence alignments and tree inference was carried out using mafft and iqtree as described above. The consensus trees were edited and annotated using the iTOL webserver (Letunic and Bork, 2021) and Adobe Illustrator.

### 2.9 Identifying nuWTs and nuMTs

The assembled nuclear genomes were aligned to the respective *Wolbachia w*Mpe or *w*Moz (Sinha et al., 2022) genomes using nucmer v4.0.0beta2 (Marçais et al., 2018). Aligned fragments (“MuM”s ) that were separated by less than 50 bp, were combined into a single nuWT locus. For manual validation of HGT of *Wolbachia* sequences into filarial contigs, the *Wolbachia* regions were verified to be flanked by nematode-like sequences and protein coding genes from nematodes.

A similar strategy was employed for nuMT identification, with nucmer alignments against respective mitochondrial sequences, (Chung et al., 2020, Crainey et al., 2018), and blastx against nematode nuclear proteomes. The nuMT loci with more than 85% AT-content were removed to avoid spurious matches to low-complexity regions.

Comparison of nuMT or nuWT loci for conservation of their sequence and position across different genomes was based on their sequence comparisons using megablast coupled with the synteny analysis described above. The sequences for nuMT or nuWT loci were extracted for each genome and compared using megablast, and only hits with more than 90% identity over 90% of the sequences were analyzed further as candidates for conserved loci. In the next step, if these candidate loci were present on syntenic contigs, they were marked as conserved loci across strains or species. A blastx analysis of the nuWT or nuMT sequences against the respective mitochondrial or *Wolbachia* proteomes was performed to identify any intact protein coding genes.

## 3 Results

### 3.1 PacBio sequencing and *de novo* assembly generates a highly contiguous *M. perstans* genome

PacBio sequencing and *de novo* genome assembly of genomic DNA isolated from *M. perstans* isolate “Mpe-Cam-1”, together with BlobTools analysis for metagenomic binning resulted in a high-quality *M. perstans* genome. The genome assembly is comprised of 581 scaffolds with a total size of 76 Mb, with the largest scaffold greater than 1.2 Mb and a N50 size greater than 285 kb (Table 1). Based on these quality metrics, the Mpe-Cam-1 assembly ranks highly in comparison to all filarial genomes available so far (Supplementary Material Table 1). Repetitive elements account for 9.7% of the genome, of which about 27% of the repeat sequence length was comprised of low complexity and simple repeats. Long terminal repeat (LTR) retrotransposons comprised 17% of all the repeats, while 50% of the repeat content is comprised of unknown, novel repeat families (Supplementary Material Table 2). The genome has 9,460 protein coding genes, with a BUSCO score of 93.2 % (Supplementary Figure 1) when evaluated against a set of 3,131 core proteins conserved across the nematode phylum, further indicating the high quality of this genome assembly.

**Table 1.** Summary statistics of Mansonella genome assemblies.

|  | *M. perstans* | *M. perstans* | *M. ozzardi* | *M. ozzardi* |
| --- | --- | --- | --- | --- |
| Isolate Name | Mpe1-Cam-1 | Mpe-Cam-2 | Moz-Brazil-1 | Moz-Venz-1 |
| Sampling site | Cameroon | Cameroon | Brazil | Venezuela |
| Sequencing Platform | PacBio | Illumina | Illumina | Illumina |
| Assembled genome size | 76 Mb | 74.35 Mb | 75.78 Mb | 74.86 Mb |
| Number of scaffolds | 581 | 6,206 | 4,789 | 4,561 |
| Largest scaffold | 1.25 Mb | 217 kb | 317 kb | 258 kb |
| Scaffold N50 size | 285.8 kb | 29kb | 37.75 kb | 38.90 kb |
| %GC | 30.31 | 30.06 | 29.65 | 29.67 |
| % Repeats | 9.70 % | 7.77 % | 8.57% | 7.91 % |
| Predicted genes | 9,477 | 9,984 | 9,834 | 9,860 |

Another clinical isolate from Cameroon, “Mpe-Cam-2” was sequenced and assembled *de novo* based on Illumina sequencing. Although the quality metrics of this Illumina short-reads assembly are lower than the PacBio long-read Mpe-Cam-1 assembly (Table 1), the repeat content, (Supplementary Material Table 2), the number of predicted genes and their BUSCO scores are similar to Mpe-Cam-1 (Supplementary Figure 1).

### 3.2 Genome sequencing and assembly of *M. ozzardi* from South America

For *M. ozzardi*, two genome assemblies from Latin American isolates, “Moz-Brazil-1” from Brazil and “Moz-Venz-1” from Venezuela, were generated based on Illumina sequencing reads. These assembled genomes are about 76 Mb (Table 1), similar in size to the *M. perstans* genomes described above. The number of predicted genes, their BUSCO scores (Supplementary Figure 1) and repeat content (Supplementary Material Table 2) are also similar to those present in *M. perstans* genomes.

### 3.3 Analysis of genome variation across *M. perstans* and *M. ozzardi* and among multiple clinical isolates

The availability of genomes from two *Mansonella* species and multiple isolates provided a unique opportunity for analysis of whole genome sequence variation both between and within the two species. The genome-wide average nucleotide identity (gANI) scores for isolates within species are 99.5% (Table 2) in comparison to the score of 89.69% between the two *Mansonella* species.

**Table 2.**
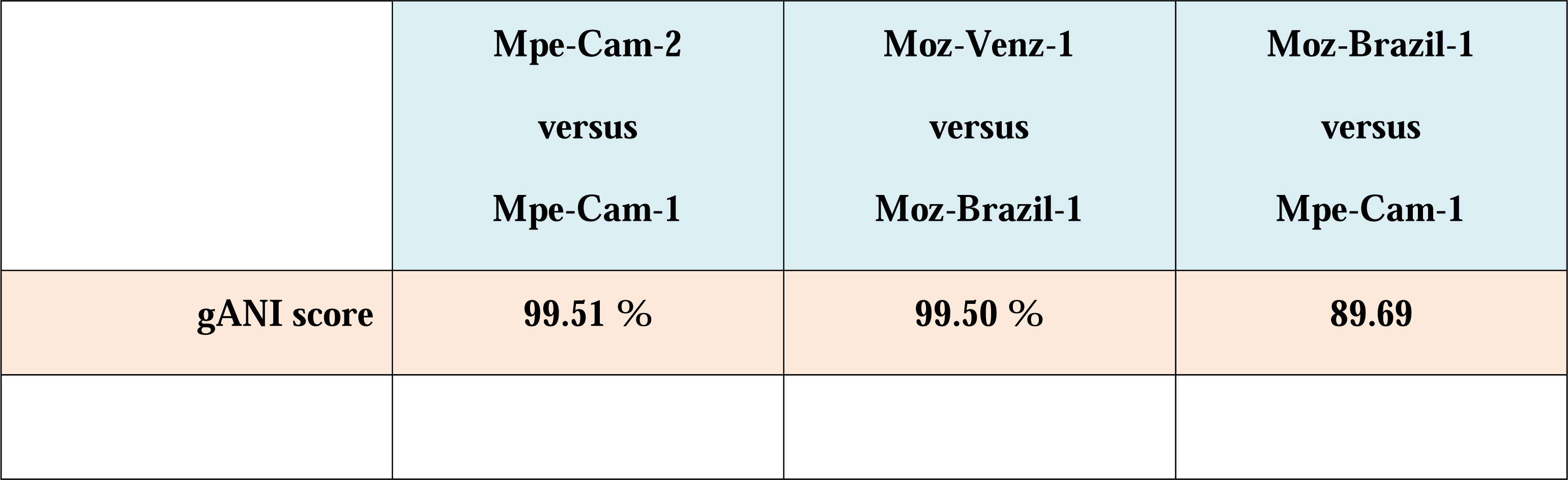
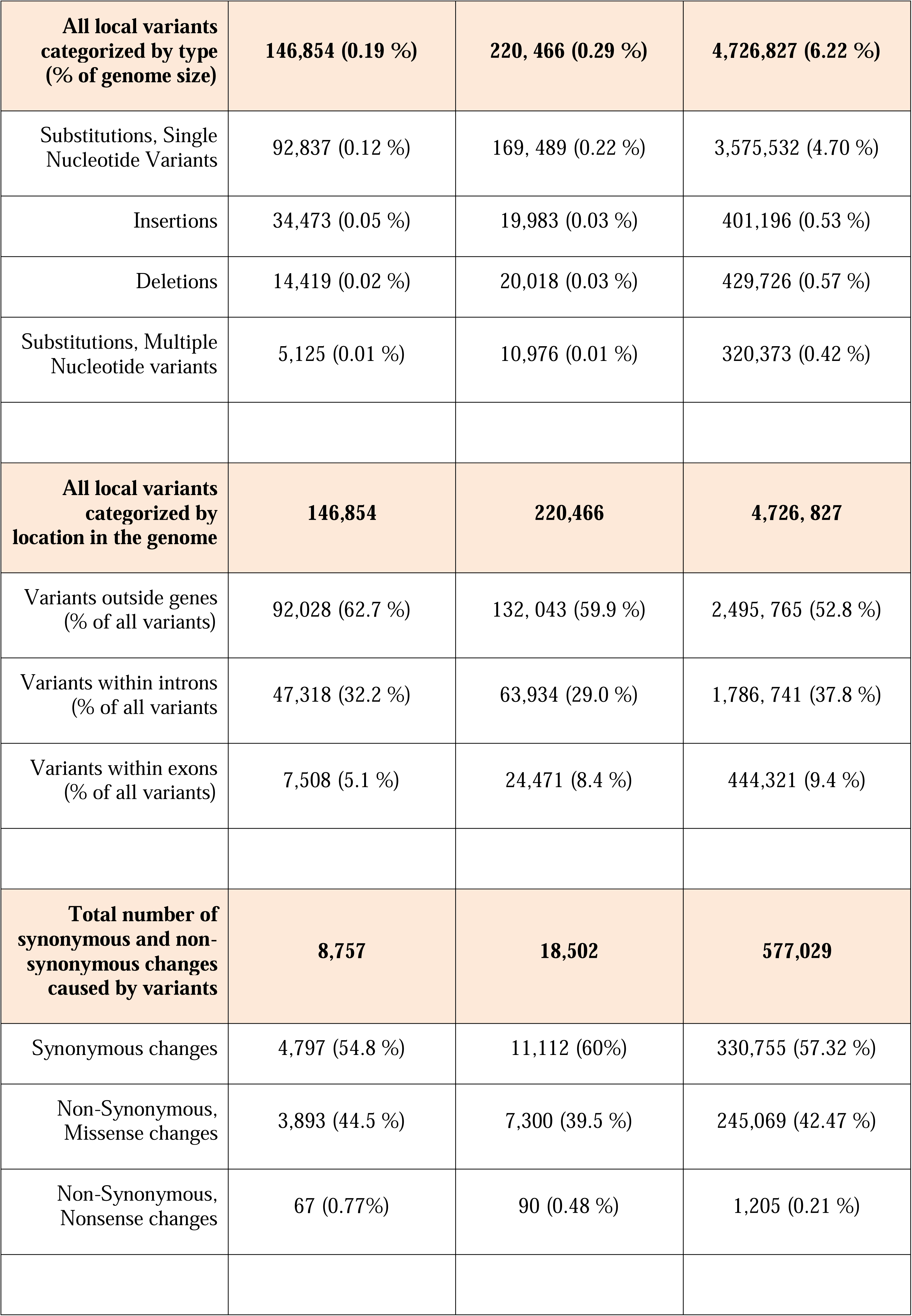
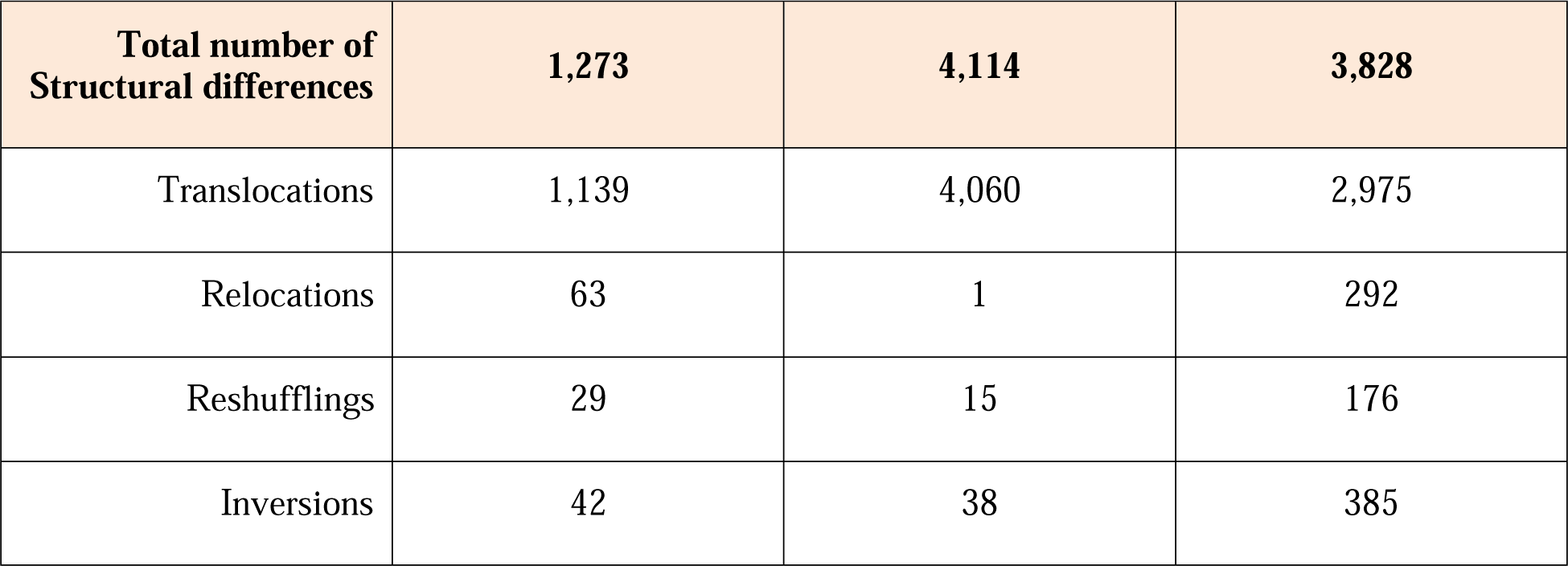
Structural and local differences between Mansonella genomes.

Visualizations of whole genome alignments between isolates as Jupiter plots (Figure 1) reveal high levels of synteny. The isolates Mpe-Cam-1and Mpe-Cam-2, collected from the same geographical region within a few years apart are most similar to each other (Figure 1A). Relatively more genomic re-arrangements (represented as crisscrossing lines in Jupiter plots) can be seen between *M. ozzardi* isolates Moz-Brazil-1 and Moz-Venz-1 (Figure 1B), which have been collected from different geographical regions (Brazil, Venezuela) and at about 30 years apart. This indicates that the evolution of *Mansonella* genomes is dependent on geographical and temporal separation. Whole-genome alignments comparing *M. perstans* and *M. ozzardi* genomes also demonstrate high levels of synteny (Figure 1C), although lower than the within-species levels, consistent with the trend in their gANI scores.

**Figure 1.**
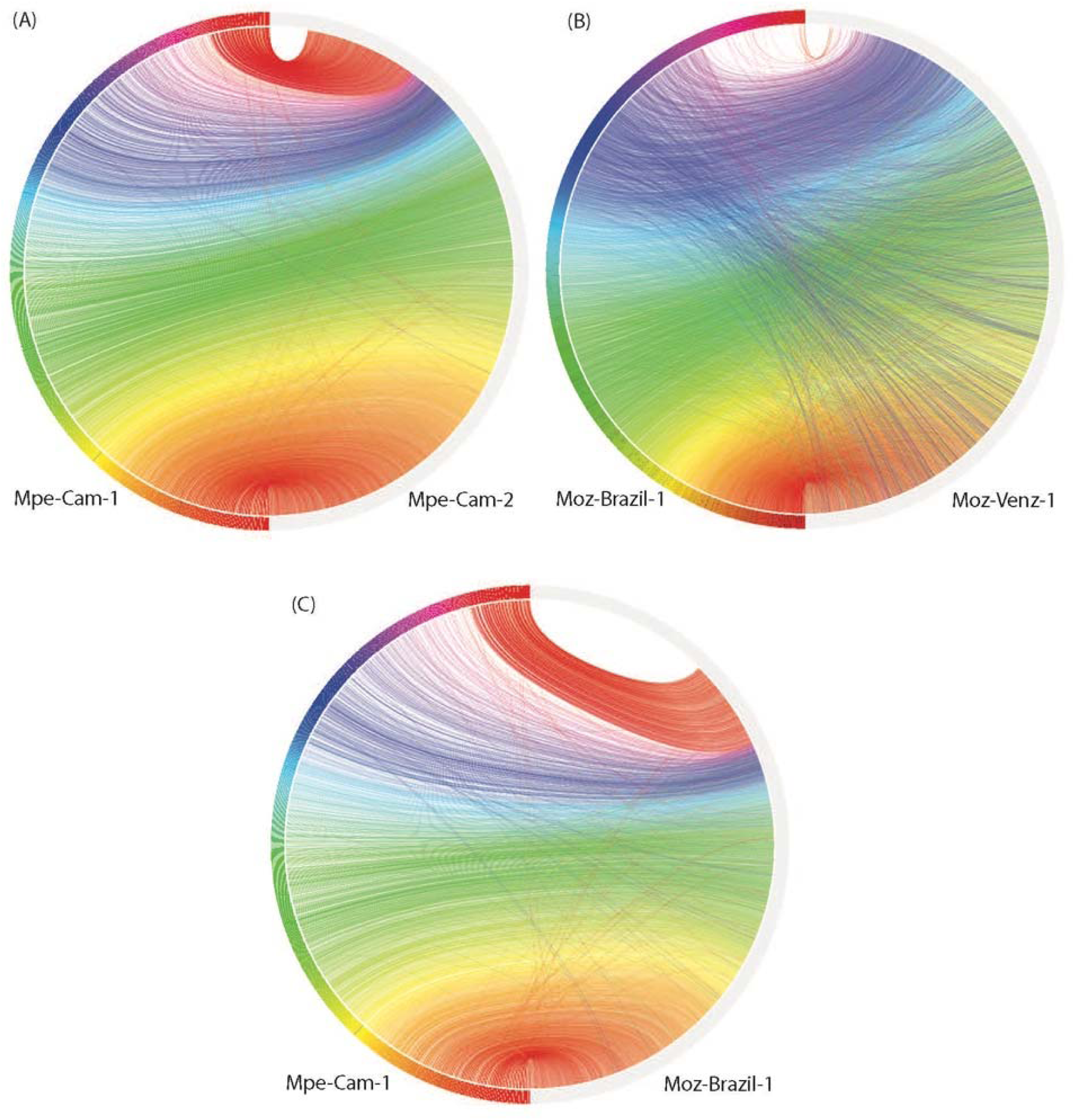
Comparisons of syntenic regions across the assembled *Mansonella* genomes. Whole genome alignments were performed using minimap with default parameters. Syntenic regions were visualized using Jupiter plots, comparing (A) The two isolates of *M. perstans* (B) The two isolates of *M. ozzardi*, and (C) the cross-species comparison between Mpe-Cam-1 and Moz-Brazil-1.

At the nucleotide level, the numbers of simple substitutions, insertions and deletions (Table 2) were 10 to 30-fold higher in across-species comparisons (6.22%), as compared to within-species (0.2% and 0.29%). In all the pairwise comparisons, most of the sequence variation (>52.8%) was localized to the intergenic portions of the genomes outside the protein-coding genes CDS regions. The proportion of variants within protein-coding genes ranged from about 5% to 10% (Table 2). Most of the nucleotide changes (54% to 60%) that occurred within coding regions, were mutationally silent as they resulted in synonymous amino-acid changes, 39 to 44% of coding variants did result in missense or non-synonymous changes in the predicted protein sequences and only a small fraction (0.21 % to 0.77%) of the coding variants resulted in a nonsense mutation. Overall, the intra-species differences between the two *M. ozzardi* isolates from different geographical areas were higher than the corresponding intra-species differences of two *M. perstans* isolates which were both collected in Cameroon. A similar trend was observed for structural differences as well, as the number of relocations, reshufflings and inversions of contig fragments was observed to be much higher in cross-species comparisons (Table 2).

### 3.4 Orthology and phylogenomic analysis of filarial nematodes

An Orthofinder analysis of proteomes from all available filarial genomes (Supplementary Material Table 1), with *S. digitata* and *T. callipaeda* as immediate outgroups from the sub-order Spiruromorpha, and *C. elegans* and *P. pacificus* as distant outgroups identified 3,829 orthogroups conserved across all these nematodes (Figure 2). This analysis also highlighted 110 orthogroups specific to *Mansonella* genus, 320 orthogroups specific to *Onchocerca* genus, and 130 orthogroups specific to *Brugia*. Most of these proteins had only weak similarities to other nematode proteins (median sequence identity and coverage both around 33%), making them good candidates for genus-specific biomarkers (Supplementary Material Table 3).

**Figure 2.**
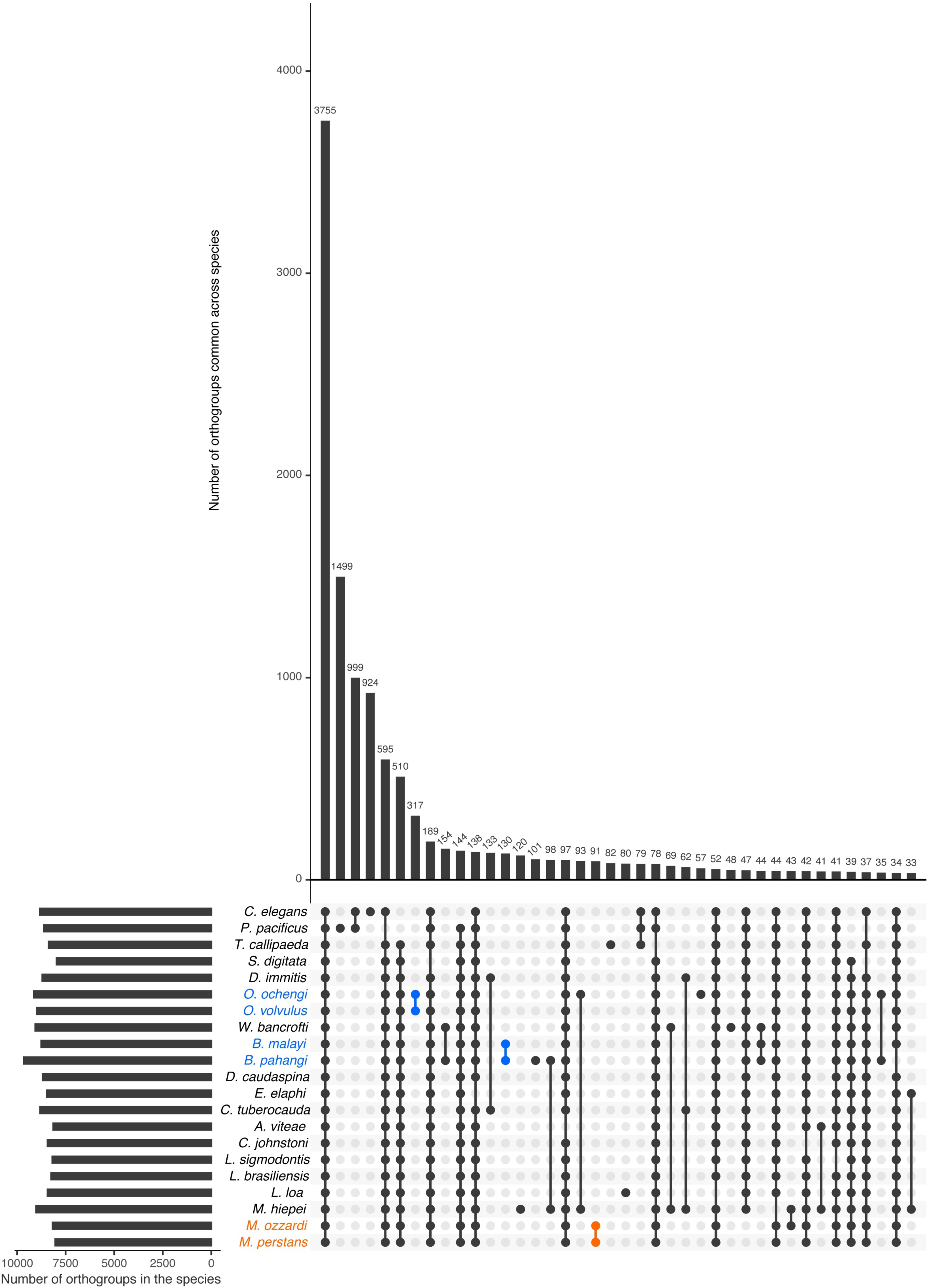
Orthology analysis across filarial parasites. Homologs conserved across various filarial species, as well as the Onchocercid nematodes *S. digitata* and *T. callipaeda*, and free-living model nematodes *P. pacificus* and *C. elegans* from Clade V were identified using OrthoFinder. The set of all 263,019 proteins from these nematode genomes were grouped into 16,564 orthogroups. The number of orthogroups identified in each species are represented by horizontal bars in the Upset plot, and the number of orthogroups common across different species are represented by vertical bars. The presence of an orthogroup in a species is indicated by a filled, colored dot while the absence is indicated by empty gray dots. Genus-specific orthogroups are shown as orange dots/bars for *Mansonella*, and blue dots/bars for *Onchocerca* and *Brugia*.

Among the orthogroups conserved across all nematode genomes analyzed here, 465 orthogroups comprised of single copy orthogroups (SCOs) and were used for analysis. The phylogenomic tree (Figure 3) generated using these 465 orthogroups recapitulated the phylogenetic relationships based on analysis of mitochondrial genomes (McCann et al., 2021) and the monophyletic feature of major clades ONC2, ONC3, ONC4 and ONC5 based on 7 genes (Lefoulon et al., 2015).

**Figure 3.**
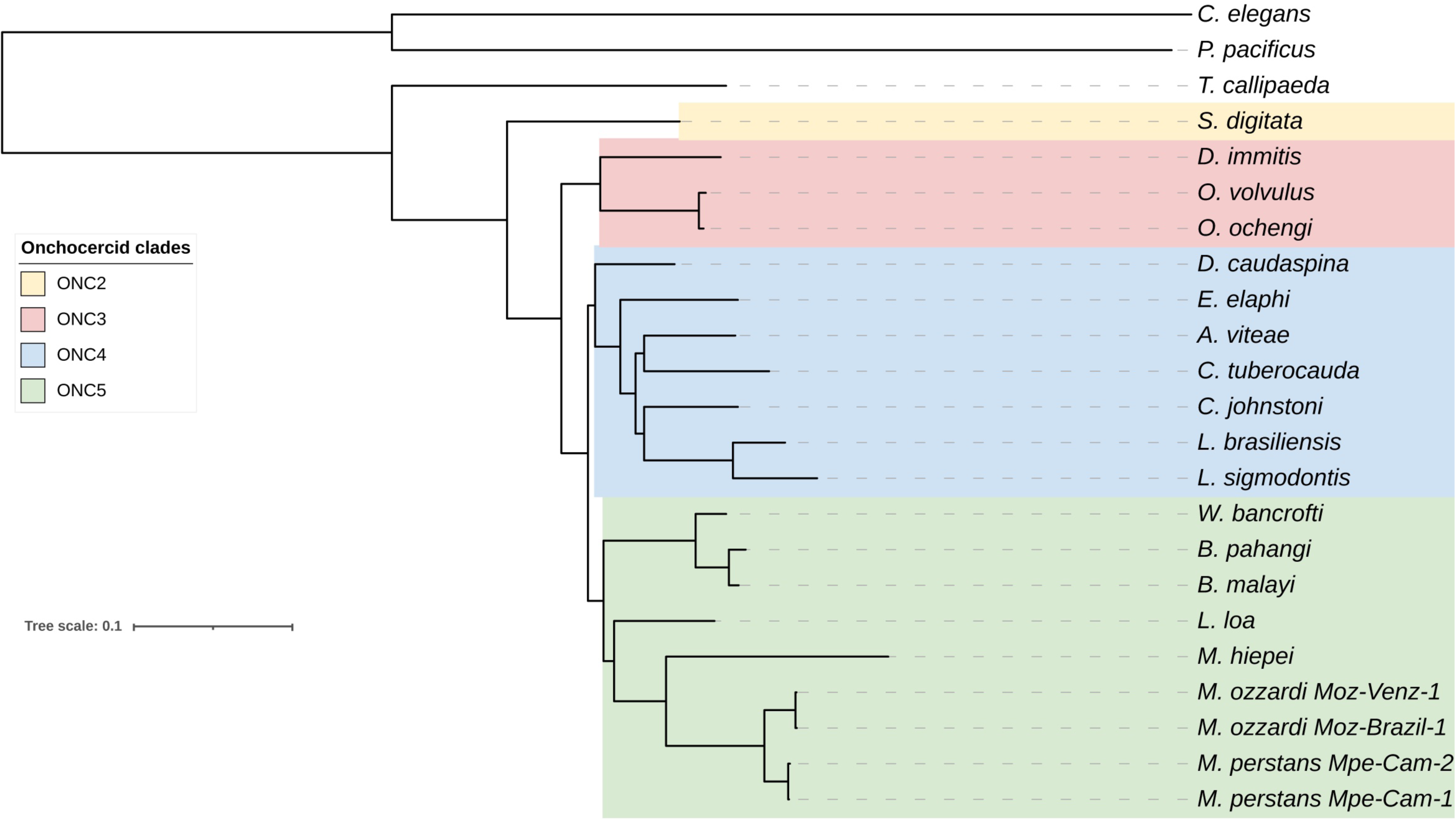
Phylogenomic analysis of filarial parasites. Phylogenomic tree was generated based on protein sequences of 465 single copy orthologs identified across all nematode genomes. Onchocercid specie *S. digitata* and *T. callipaeda* were used as immediate outgroups, while free-living nematodes *C. elegans* and *P. pacificus* were used as more distant outgroups. The previously described Onchocercid clades ONC2 to ONC5 are highlighted.

### 3.5 Analysis of nuMTs, horizontal transfers of mitochondrial DNA into nuclear genomes

Transfer of mitochondrial DNA fragments into the nuclear genome has been observed for multiple filarial nematodes (Grote et al., 2017) as well other eukaryotes (Hazkani-Covo et al., 2010). Since mitochondrial markers are often used as barcodes for species identification and phylogenetic analysis (Crainey et al., 2018), it is important to distinguish between genuine mitochondrial sequences versus nuMTs (Song et al., 2008, Crainey et al., 2018). In the newly sequenced *Mansonella* genomes, 39 to 94 nuMT loci with median sizes ranging around 250 bp were found across different isolates, with the largest loci ranging from 2.35 kb to 3.69 kb in size (Table 3). None of the nuMTs was found to carry an intact protein coding gene. The total number and the combined size of nuMTs in was observed to be higher in *M. ozzardi* as compared to *M. perstans* (Table 3). The locations and sequences of nuMTs were poorly conserved across *M. perstans* and *M. ozzardi*. For example, in a comparison of the 48 Mpe-Cam-1 nuMTs with the 82 Moz-Brazil-1 nuMTs, only 2 conserved loci were found (Supplementary Material Table 4). Even within the isolates of each species, approximately 35% and 60% of nuMT loci were conserved in *M. perstans* and *M. ozzardi* respectively (Supplementary Material Table 4), while the remainder were isolate-specific, indicating a fast turnover of these elements. The existence of multiple cox1 derived nuMT pseudogenes has been reported in *M. ozzardi* (Crainey et al., 2018). Similarly, multiple fragments of the *cox1* gene were also observed in both *M. ozzardi* isolates (Supplementary Figure 2). However, an exact match for these sequences could not be found in either of the Moz-Brazil-1 or Moz-Venz-1 assemblies, providing further evidence for poor conservation of such nuMTs across different isolates.

**Table 3.** Nuclear mitochondria Transfers (nuMTs) annotated across all genomes.

| Species | Isolate | Total number of nuMTs | Total size of nuMTs | Median size of nuMTs | Largest nuMT | Median percent identity to mitochondria |
| --- | --- | --- | --- | --- | --- | --- |
| *M. perstans* | Mpe-Cam1 | 48 | 19.240 kb | 238 bp | 2,591 bp | 87.81 % |
| *M. perstans* | Mpe-Cam2 | 39 | 14.330 kb | 249 bp | 2,351 bp | 86.48 % |
| *M. ozzardi* | Moz-Brazil-1 | 82 | 35.070 kb | 271 bp | 3,170 bp | 89.57 % |
| *M. ozzardi* | Moz-Venz-1 | 94 | 35.851 kb | 223 bp | 3,697 bp | 89.50 % |

**Table 4.** Nuclear Wolbachia Transfers (nuWTs) annotated across all genomes.

| Species | Isolate | Total number of nuWTs | Total size of nuWTs | Median size of nuWTs | Largest nuWT | Median percent identity to *Wolbachia* |
| --- | --- | --- | --- | --- | --- | --- |
| *M. perstans* | Mpe-Cam1 | 328 | 87.83 kb | 173 bp | 2,733 bp | 91.51 % |
| *M. perstans* | Mpe-Cam2 | 340 | 76.26 kb | 171 bp | 1,236 bp | 91.84 % |
| *M. ozzardi* | Moz-Brazil-1 | 308 | 77.33 kb | 198 bp | 1,515 bp | 91.37 % |
| *M. ozzardi* | Moz-Venz-1 | 294 | 73.54 kb | 195 bp | 2,151 bp | 90.97 % |

When the nuMT sequences from each isolate were mapped back to the mitochondrial genomes, they were found to cover almost the entire mitochondrial genome, without any preference for specific regions to be transferred into the nuclear genome (Supplementary Figures 2 and 3).

### 3.6 Analysis of nuWTs, horizontal transfers of *Wolbachia* endosymbiont DNA into host nuclear genomes

Instances of transfer of DNA sequences from *Wolbachia* endosymbionts into the nuclear genomes of their hosts, termed “nuWTs” for nuclear-*Wolbachia* transfers, have been observed frequently in filarial nematodes (McNulty et al., 2010; Cotton et al., 2016; Grote et al., 2017; Hotopp et al., 2017) as well as arthropod hosts (Klasson et al., 2014; Tvedte et al., 2022). The *Mansonella* isolates reported here harbor about 300 nuWT loci in each isolate, with median size about 180 bp, adding up to a total of 76 kb to 88 kb), with the largest nuWT (2.73 kb) observed in Mpe-Cam-1 (Table 4). None of these nuWTs were found to harbor any intact protein coding gene. Analysis of synteny and sequence identity of nuWTs across species identified only 34 conserved nuWTs between *M. perstans* and *M. ozzardi* (Supplementary Material Table 5). Interestingly, about 80% (260 from *M. perstans* and 280 from *M. ozzardi*) were found to be conserved in within-species comparisons of *M. perstans* and *M. ozzardi* respectively (Supplementary Material Table 5).

### 3.7 Analysis and comparison of genes encoding drug targets for the filaricides diethylcarbamazine, ivermectin and albendazole

The main preventive intervention for filarial infections is based on mass drug administration (MDA) using diethylcarbamazine (DEC), ivermectin (IVM) and albendazole (ALB) (Simonsen et al 2010). Although DEC has been successfully used to treat *M. perstans* infections, the drug has been found to be ineffective against *M. ozzardi* (Chadee et al., 1995). To investigate potential reasons for differences in drug susceptibility between *M. perstans* and *M. ozzardi*, the genes encoding the drug targets for DEC, ivermectin and albendazole, were identified and analyzed. A search for the DEC target *gon-2* (Verma et al., 2020), a gene that encodes a Transient Receptor Protein M (TRPM) channel subunit, identified two potential homologs in *M. perstans* but one in *M. ozzardi*. Phylogenetic analysis of the *gon-2* homologs in *Mansonella* and other nematode genomes revealed two distinct, paralogous clades, denoted as Clade 1 and Clade 2, which represent filarial orthologs of *Cel-gon-2* and *Cel-gtl-2* respectively (Figure 4). The nematodes known to be susceptible to DEC, including *C. elegans* and filarial parasites *B. malayi L. loa*, and *W. bancrofti*, all have a *gon-2* ortholog in Clade 1. Interestingly, *M. perstans*, which is treatable with DEC, also has a corresponding ortholog in Clade 1 while in *M. ozzardi*, which is refractory to DEC, the orthologous gene was found in two fragments on the ends of two separate contigs in each of the two isolates (Supplementary Figure 4). Artificially combining the two gene fragments could reconstruct the full gene except for a few missing bases between exons 19 and 20 (Supplementary Figures 5 and 6). It is possible that the gene is intact in *M. ozzardi* but is split over two contigs due to incomplete genome assembly. To explore this further, the raw Illumina reads from each isolate were mapped to a synthetic contig constructed by joining the contigs carrying the *gon-2* fragments in appropriate orientations with a 50 bp spacer marked by Ns, and all read-pairs spanning the potentially un-assembled gap were investigated (Supplementary Figures 7 and 8). This analysis showed an abrupt drop in read coverage in the potential gap region, even though the contig terminals had sufficient coverage. Although a few read-pairs (less than 10 counts) were observed to span across the gap, the size of gap estimated via the insert size of the paired-end reads was around 2.5 kb, much larger than the insert size of the sequencing library. In contrast, the read-coverage across the corresponding region in *M. perstans* is uniform (Supplementary Figure 9). Together, these observations indicate that the synthetic contig is unlikely to represent the true genomic sequence, and therefore the *gon-2* gene is fragmented in both *M. ozzardi* isolates. Both *M. perstans* and *M. ozzardi* do possess orthologs of *Cel-gtl-2* in Clade 2, which share only 52 % sequence identity with the *C. elegans* GON-2 protein in Clade 1 and are unlikely to share their functions. Similar analysis for the other known DEC target genes, *trp-2*, *ced-11* and *slo-*1 identified the corresponding one-to-one orthologs for all three genes in all the filarial species including both isolates of *M. perstans* and *M. ozzardi* (Supplementary Figures 10, 11 and 12). Taken together, this suggests that the absence the *gon-2* ortholog in *M. ozzardi* is most likely responsible for the ineffectiveness of DEC against this parasite.

**Figure 4.**
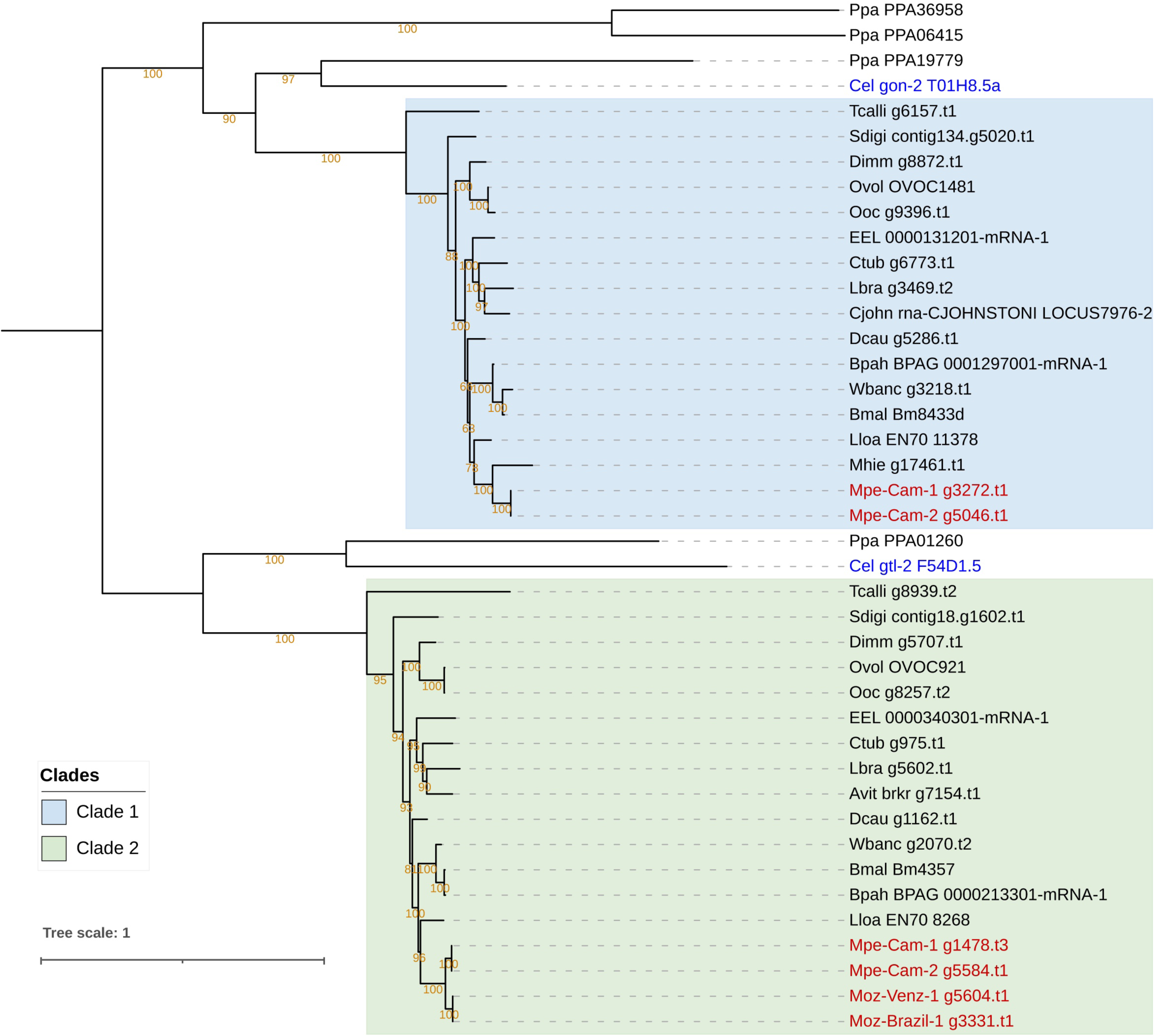
Phylogenetic analysis of genes encoding the targets of the anti-filarial drug Diethylcarbamazine (DEC). Phylogenetic analysis of the *gon-2*, *gtl-1* and *gtl-2* gene family was performed based on their protein sequences and the tree was rooted at the midpoint. The orthologs of the *Cel-gon-2* gene are placed in Clade 1, highlighted in blue, and orthologs of *Cel-gtl-2* orthologs are placed in Clade 2, highlighted green. The *gtl-1* gene has no orthologs in filarial nematodes. Genes from *M. perstans* and *M. ozzardi* are in red, and *C. elegans* genes are in blue. Asterisks (*) next to gene names indicate fragmented gene structures.

The targets of ivermectin as identified through genetic studies in *C. elegans* include various subunits of the GlutamateLgated Cl− channels (GluCl), encoded by genes *glc-1*, *glc-2*, *glc-3*, *glc-4*, *avr-14* and *avr-15* (Dent et al., 2000; Wolstenholme and Rogers, 2005). Ivermectin is effective in treating *M. ozzardi* infections (Basano et al., 2018), but is reported to be ineffective against *M. perstans* (Bregani et al., 2006). Phylogenetic analysis of these genes across filarial parasites shows that the homologs of *glc-1*, *glc-2*, *glc-3*, *avr-14* and *avr-15* form a family of paralogous genes with complex patterns of homology across filarial parasites (Figure 5), while *glc-4* has one-to-one orthologs across all genomes analyzed (Supplementary Figure 13). Orthologs for *Cle-glc-1*, *Cel-glc-2* and *Cel-avr-15* are not present in either of the *Mansonella* species (Figure 5). Both isolates of *M. perstans* and *M. ozzardi* have one ortholog most closely related to the *Cel-glc-3*, and one ortholog most closely related to to *Cel-avr-14*,

**Figure 5.**
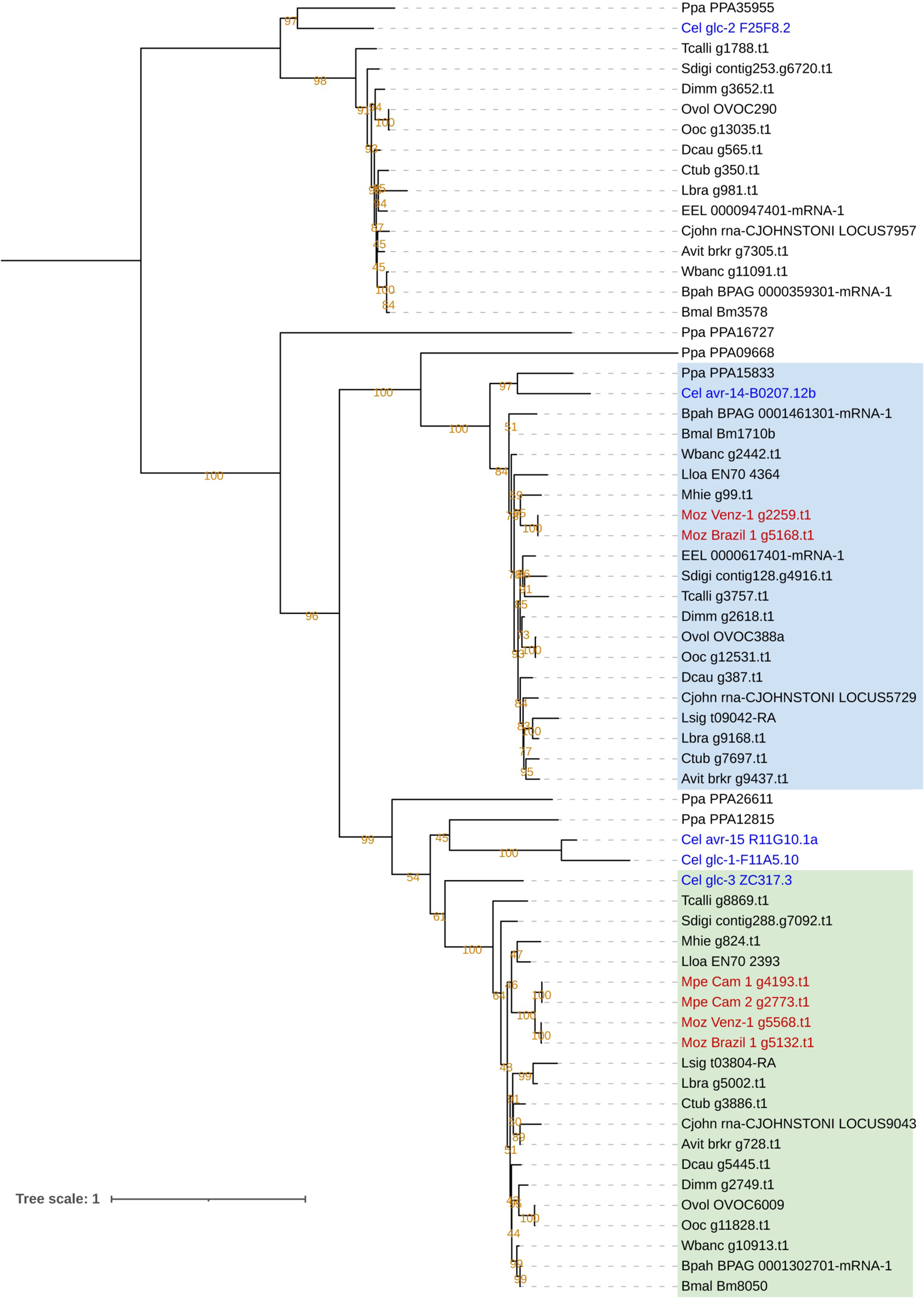
Phylogenetic analysis of genes encoding Glutamate Chloride channel subunits, targets of ivermectin. Phylogenetic analysis of the GluCl gene family was based on their protein sequences and the tree was rooted at the midpoint. The clade with *Cel-glc-2* orthologs is highlighted in yellow, clade with *Cel-avr-14* orthologs is highlighted in blue, and the clade with *Cel-glc-3* orthologs is highlighted in green. *M. perstans* and *M. ozzardi* genes are in red, and *C. elegans* genes are in blue.

The anthelminthic drug mebendazole acts by binding to beta-tubulin protein and inhibiting microtubule formation (Lacey, 1990). Mebendazole is commonly used in combination DEC or ivermectin for treating *M. perstans* infections (Bregani et al., 2006) but its use for *M. ozzardi* has not been reported. The beta-tubulin gene family in *C. elegans* consists of 6 different members (*ben-1*, *tbb-1*, *tbb-2*, *tbb-4*, *tbb-6*, and *mec-7*) all of which have varying effects on drug sensitivity (Driscoll et al., 1989; Pallotto et al., 2022). A phylogenetic analysis of the beta-tubulin gene family in *Mansonella* and other nematodes reveals a complex pattern of expansion and contraction of this gene family across filarial species, with no direct orthologs for *ben-1* identifiable in any filarial nematodes (Supplementary Figure 14). The exact roles of different beta-tubulin homologs on drug susceptibility in *Mansonella* will require further study.

## 4 Discussion

To gain some insight into the neglected pathogens responsible for human mansonellosis, we report the first *de novo* genome assemblies from the filarial parasites *M. perstans* and *M. ozzardi*, providing reference genomes for each species. The genomes encode approximately 10,000 genes each and are high quality and largely complete based on universally conserved orthologs (BUSCO scores greater than 90%) relative to other filarial species. The availability and sequencing of two isolates for *M. perstans* and *M. ozzardi* also allowed higher confidence in calling presence or absence of genes in each species as it could be confirmed in two independent isolates. The *Mansonella* genomes average 76 Mb in length and are similar in size to the genomes described from other human and animal filarial parasites (Ghedin et al., 2007; Godel et al., 2012; Desjardins et al., 2013; Tallon et al., 2014; Lau et al., 2015; Foster et al., 2020, McCann et al., 2021). The sequencing of two clinical isolates of *M. perstans* and *M. ozzardi* also enabled a quantification of the genetic variability both between species as well in the same parasite from different regions. Most of the genetic differences observed between isolates and species are in the intergenic regions, and only a small fraction (10%) of variants affects coding exons. Relatively little is known about natural variation in the nuclear genomes of filarial parasites (McNulty et al 2013., Qing et al., 2021). In the heartworm *Dirofilaria immitis*, a closely related filarial parasite of dogs and cats, sequencing of two isolates from Europe and the United States revealed very little genetic variation in their nuclear or mitochondrial genomes (Godel et al., 2012). Other studies of genetic variation within isolates of a filarial species have used mitochondrial *cox1* gene (McNulty et al., 2013) or the mitochondrial genomes (Qing et al., 2021). Our study presents a more comprehensive analysis of variation in nuclear genomes within isolates. It will be interesting to investigate how these patterns evolve in other human filarial species.

Transfer of mitochondrial DNA fragments into the nuclear genome has been observed for multiple filarial nematodes (Grote et al., 2017). These nuMT loci are annotated for all *Mansonella* isolates using their reported mitochondrial genomes (Crainey et al., 2018; Chung et al., 2020). Most filarial parasites possess the intracellular bacterial symbiont, *Wolbachia*, capable of transferring genetic material to the nuclear genome of its host. The genomes of *Wolbachia* endosymbionts *w*Mpe and *w*Moz from *M. perstans* and *M. ozzardi* have also been reported (Sinha et al., 2022), enabling an analysis of nuWTs, regions of DNA transfers from *Wolbachia* to its host nuclear genomes. While reported in other filarial species, the biological function and significance of such nuclear transfer events are not well understood. Moreover, the levels of conservation of such loci across multiple isolates of a species, or closely related species has not been previously studied. Our analysis combining synteny and sequence identity shows that the nuMT loci are poorly conserved in *Mansonella* even within the same species, and rarely harbor intact coding genes, suggesting that most of these transfers are not evolutionarily selected, and non-functional. For the nuWTs, relatively more loci were found to be conserved across the two species, but isolate specific acquisition of these nuWT fragments was also observed. However, none of these loci harbored any intact protein-coding genes, suggesting a lack of functional constraints on their evolution. This observation is consistent with the detection of mostly non-functional nuWTs in other filarial parasites (Grote et al., 2017) and suggests that these loci most likely arise randomly and degrade via genetic drift over shorter time scales, as seen in other organisms (Tvedte et al., 2022; Hazkani-Covo et al., 2010; Bensasson et al. 2003).

Despite their genetic relatedness, *M. perstans* and *M. ozzardi* differ in their geographical distribution, adaptation to insect vectors, and in their response to drug treatments. We have used these new genomes to explore the genetic basis for the reported differences in their response to commonly used anti-filarial drugs. Our analysis of the sequences and phylogeny of known gene products targeted by DEC revealed that *M. ozzardi* does not have an intact *gon-2* gene, one of the targets of this drug. In contrast, *M. perstans* as well as other filarial parasites that are susceptible to DEC have a *gon-2* ortholog. It can therefore be hypothesized that the absence of *gon-2* ortholog in *M. ozzardi* has rendered it unsusceptible to DEC. Future research focusing on this gene sequence and its variation between different isolates of the two species can shed further light on its function.

All the genes known to be targets of ivermectin were found in both *M. perstans* and *M. ozzardi*. The observed differences in ivermectin susceptibility of the two parasites could not be readily explained by the subtle differences in sequences of these targets. There could be additional ivermectin targets that are not discovered yet but could play a role in the differential response of *M. perstans* and *M. ozzardi*. Although mebendazole is used for the treatment *M. perstans*, it was found to lack an ortholog of one of the targets encoded by the beta-tubulin gene *tbb-4*, while the corresponding ortholog is present in *M. ozzardi*. Given the complex patterns of homology for this gene family, and the absence of any reported effects of mebendazole on *M. ozzardi*, the significance of this observation remains to be determined. In addition to these observations, there could be yet other undiscovered filarial genes that have also have effects on drug susceptibility and could be responsible for differences between *M. perstans* and *M. ozzardi*. A functional validation of these hypotheses regarding drug and their target genes is currently not feasible, but future advances may be able to test these hypotheses. At least one of the parasites, *M. perstans*, can now be successfully cultured *in vitro* enabling functional investigations into its biology (Njouendou et al., 2019). Advances in genome editing techniques for non-model organisms including helminth parasites open further possibilities of genetic and functional studies, and control of parasites (Lok et al., 2016; Zamanian and Andersen, 2016; Liu et al., 2020; Kwarteng et al., 2021; Sankaranarayanan et al., 2021; Quinzo et al., 2022). The availability of the new *Mansonella* genomes reported here will be a critical genomic resource for such endeavors and for further studies on drug targets, as well as enabling insights into the biology and evolution of these parasites.

## 5 Conflict of Interest

Authors AS, ZL, CBP, LE, RDM, and CKSC are employees at New England Biolabs Inc., Ipswich, MA 01938 (USA), which manufactures biotechnology reagents including those used in Next Generation Sequencing. The authors declare that the research was conducted in the absence of any commercial or financial relationships that could be construed as a potential conflict of interest.

## 6 Author Contributions

NFL, MUF, FFF, SW collected the clinical samples. AS, ZL, CBP, LE, RDM performed the sequencing experiments and bioinformatics analysis. AS, ZL, CBP, LE, RDM, CKSC analysed the data. AS, ZL, CBP, CKSC wrote the initial manuscript. AS, ZL, CBP, LE, RDM, MUF, SW, CKSC and all authors contributed to the final manuscript.

## 7 Funding

Research funding was provided by New England Biolabs, Ipswich, MA 01938, USA to AS, ZL, LE, RDM, CBP, CKSC. FAPESP (Fundação de Amparo à Pesquisa do Estado de São Paulo) supported the field research in Brazil through a research grant to M.U.F (2013/12723-7) and a doctoral scholarship to N.F.L. (2013/ 26928-0).

## 8 Acknowledgments

The authors thank the DNA sequencing core at New England Biolabs. The authors are grateful to the late Don Comb for initiating and inspiring research into NTDs at NEB.

## 9 Data Availability Statement

Raw sequencing reads for the *M. per*stans isolates Mpe-Cam-1 and Mpe-Cam-2 have been deposited in the NCBI SRA database under BioProject accession numbers PRJNA917716 and PRJNA917721 respectively. Illumina sequencing data for *M. ozzardi* isolates Moz-Brazil-1 and Moz-Venz-1 have been deposited in the NCBI SRA database under BioProject accession numbers PRJNA917722 and PRJNA917766 respectively.

## Notes

### Summary of Updates

The gene-prediction pipeline was revised to correct for gene-fusion errors and run-to-run differences in braker2 output. The avr-14 gene was found to be present in M. perstans and M. ozzardi. The gon-2 gene was present in M. perstans, while fragments were found in M. ozzardi, at the ends of two sperate contigs.

