## Supplementary Figures 1 to 14 for "Genomes of the human filarial parasites *Mansonella perstans* and *Mansonella ozzardi*"

### Supplementary Material

#### 1.1 Supplementary Figure 1. BUSCO scores of filarial parasite genomes

The completeness of the protein-coding genes in various nematode genomes was assessed using the BUSCO pipeline v5.0 beta using the “nematoda_odb10” reference dataset which has 3,131 single copy orthologs conserved across most nematode genomes.

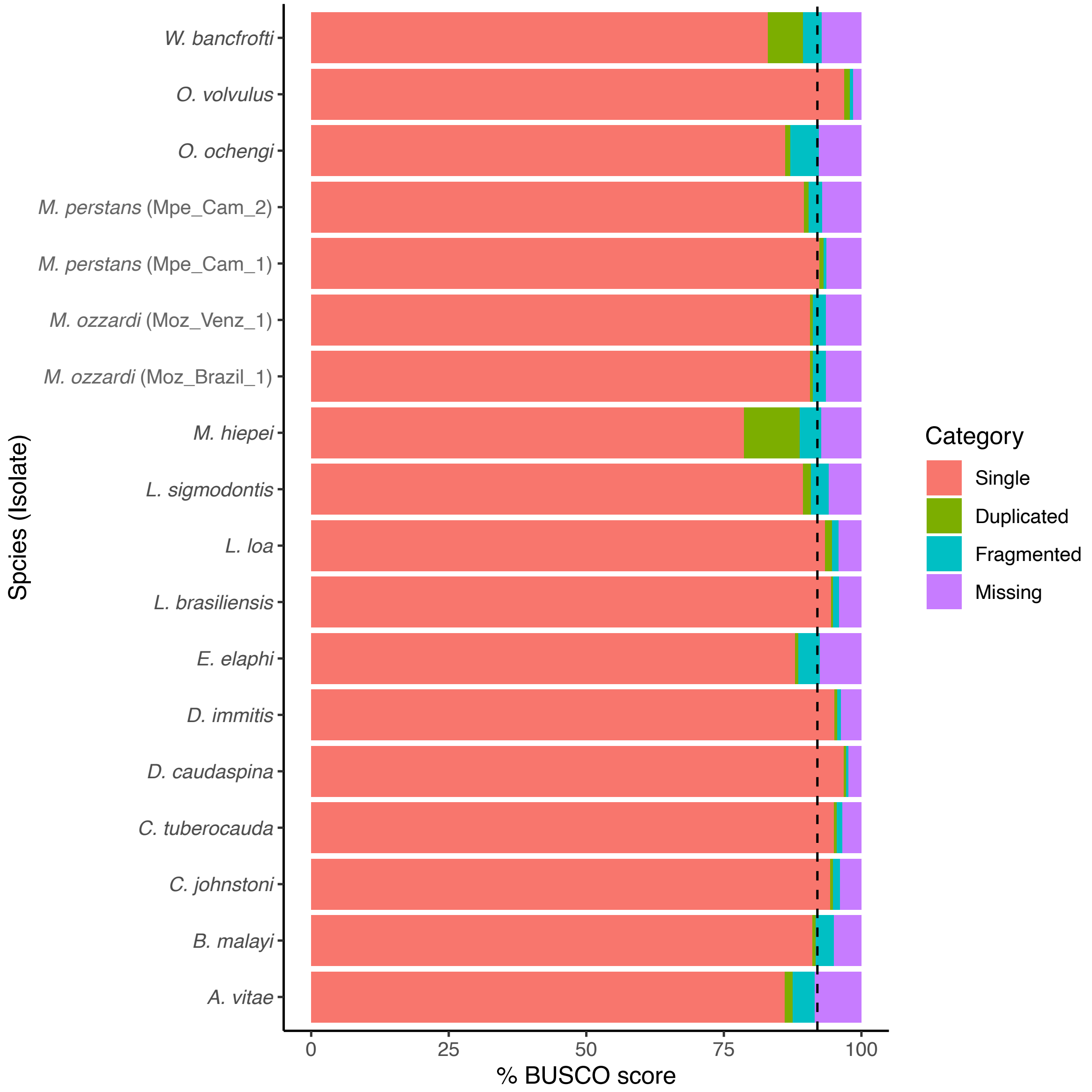

#### *1.2* Supplementary Figure 2. Source of nuMTs in *M. perstans*

The nuMT loci from each *M. perstans* isolate was mapped back to the reference mitochondrial genome of *M. perstans*. Each nuMT is represented as a horizontal bar, colored according to the percentage sequence identity of the nuMT versus the reference mitochondria.

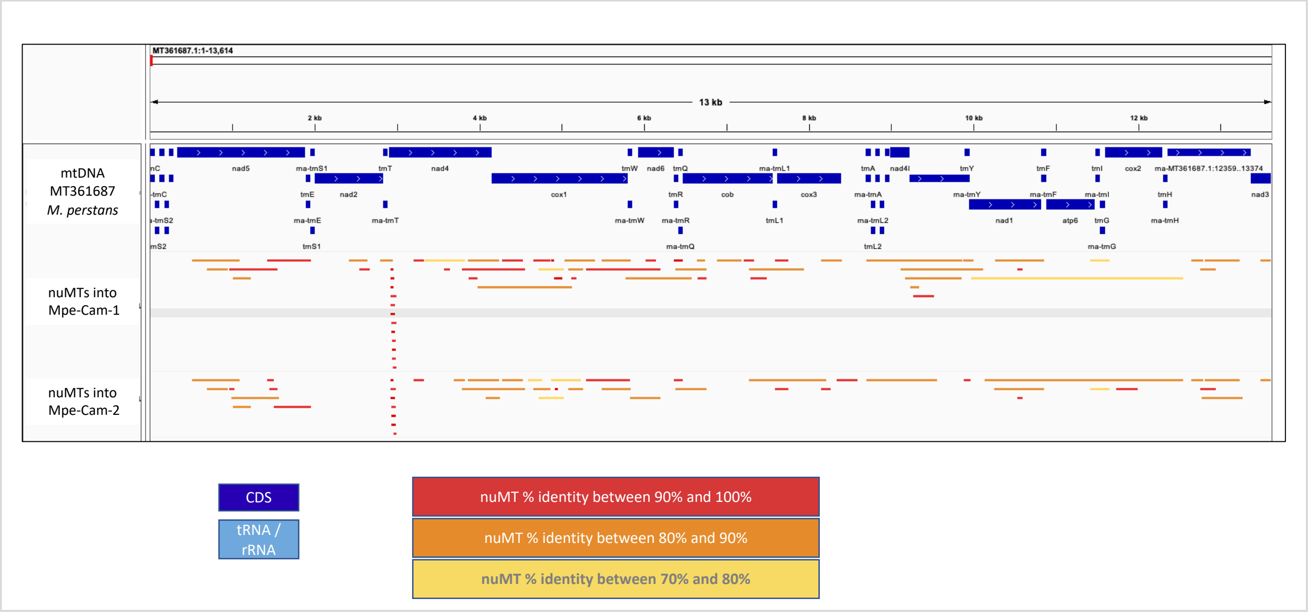

#### 1.3 Supplementary Figure 3. Source of nuMTs in *M. ozzardi*

The nuMT loci from each *M. ozzardi* isolate was mapped back to the reference mitochondrial genome of *M. perstans*. Each nuMT is represented as a horizontal bar, colored according to the percentage sequence identity of the nuMT versus the reference mitochondria.

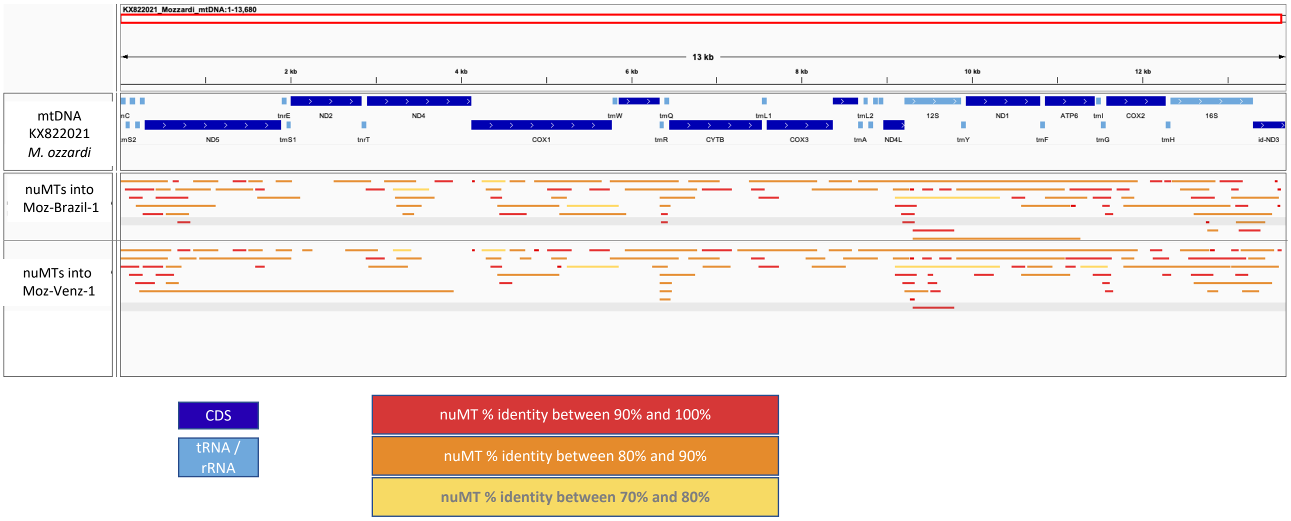

#### 1.4 Supplementary Figure 4. The *gon-2* gene is present as two fragments on two different contigs in both *M. ozzardi* isolates.

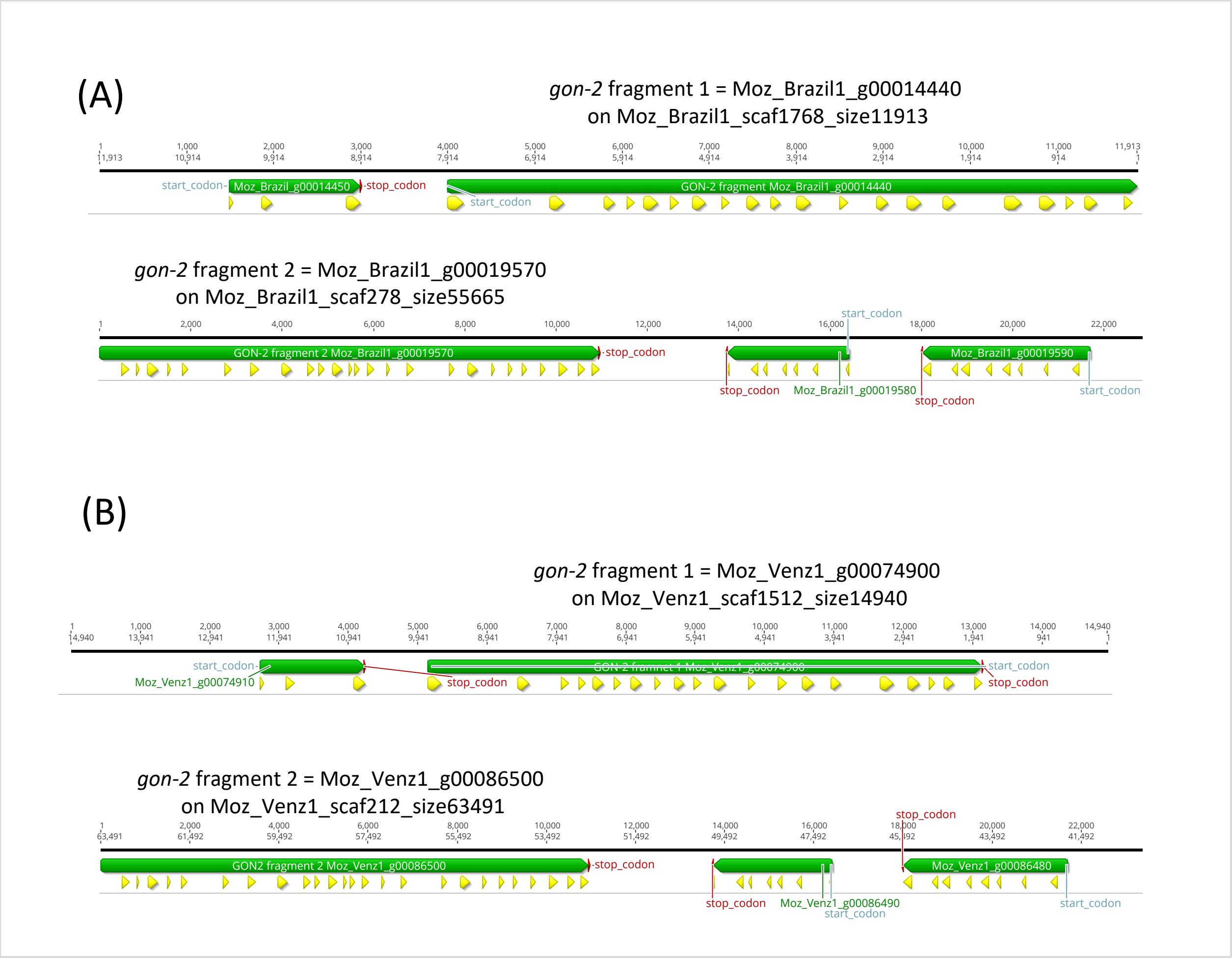

#### 1.5 Supplementary Figure 5. Pfam domain annotations for the *Cel-gon-2* gene, the corresponding orthologs in *M. perstans*, and the fragments of the *gon-2* gene in *M. ozzardi*.

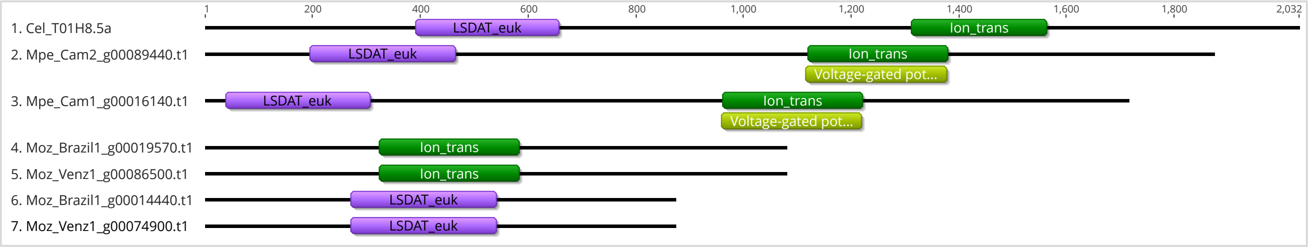

#### 1.6 Supplementary Figure 6. Alignments of the *gon-2* fragments in the two *M. ozzardi* isolates against the intact gene in Mpe-Cam-2

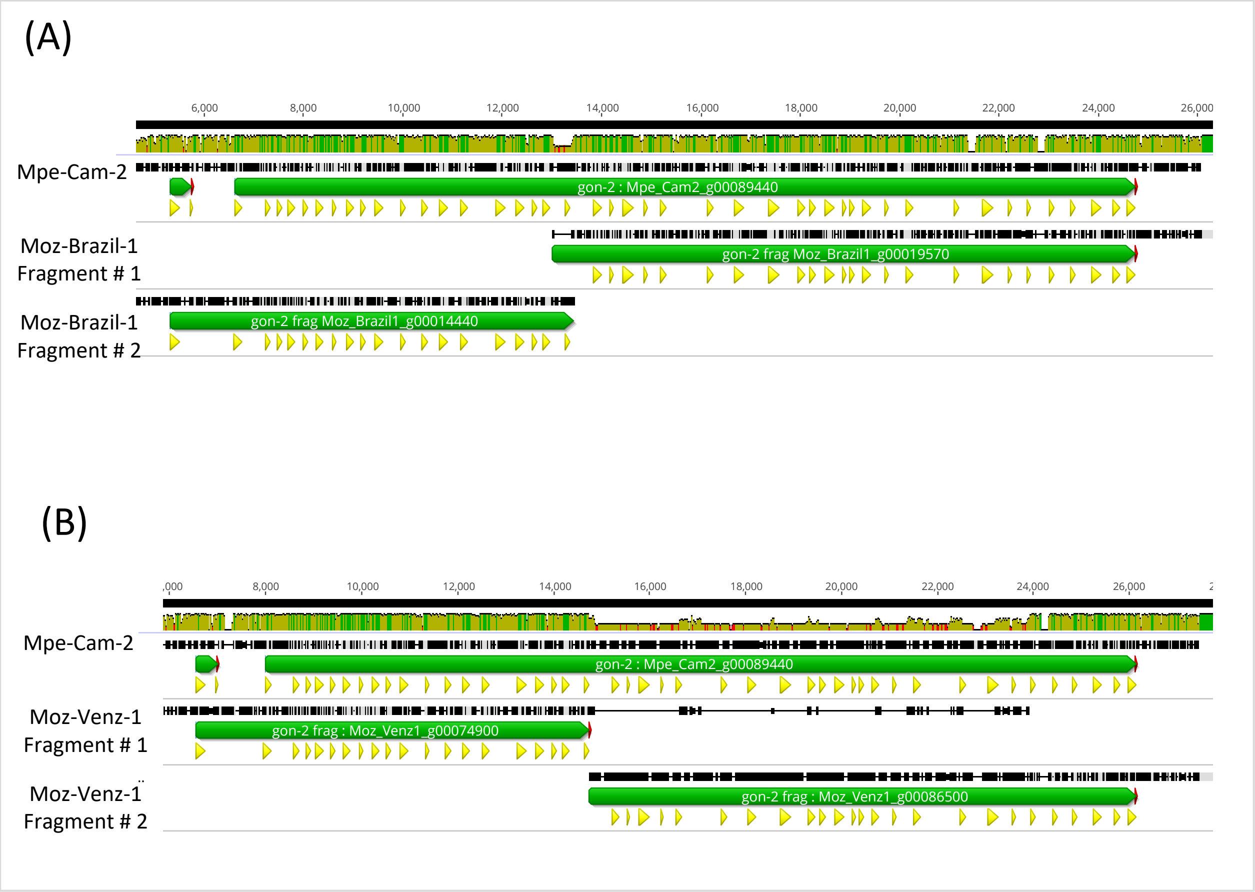

#### 1.7 Supplementary Figure 7. Read coverage analysis of the two contigs carrying the *gon-2* gene fragments in *M. ozzardi* isolate Moz-Brazil-1.

The two contigs were tentatively joined in the orientation that would reconstruct an intact *gon-2* gene. The two contigs were separated by a stretch of 50 Ns, and Illumina reads used for assembly were mapped to the synthetic contig. The bottom track shows all reads. The top track shows only the subset of reads mapped as pairs.

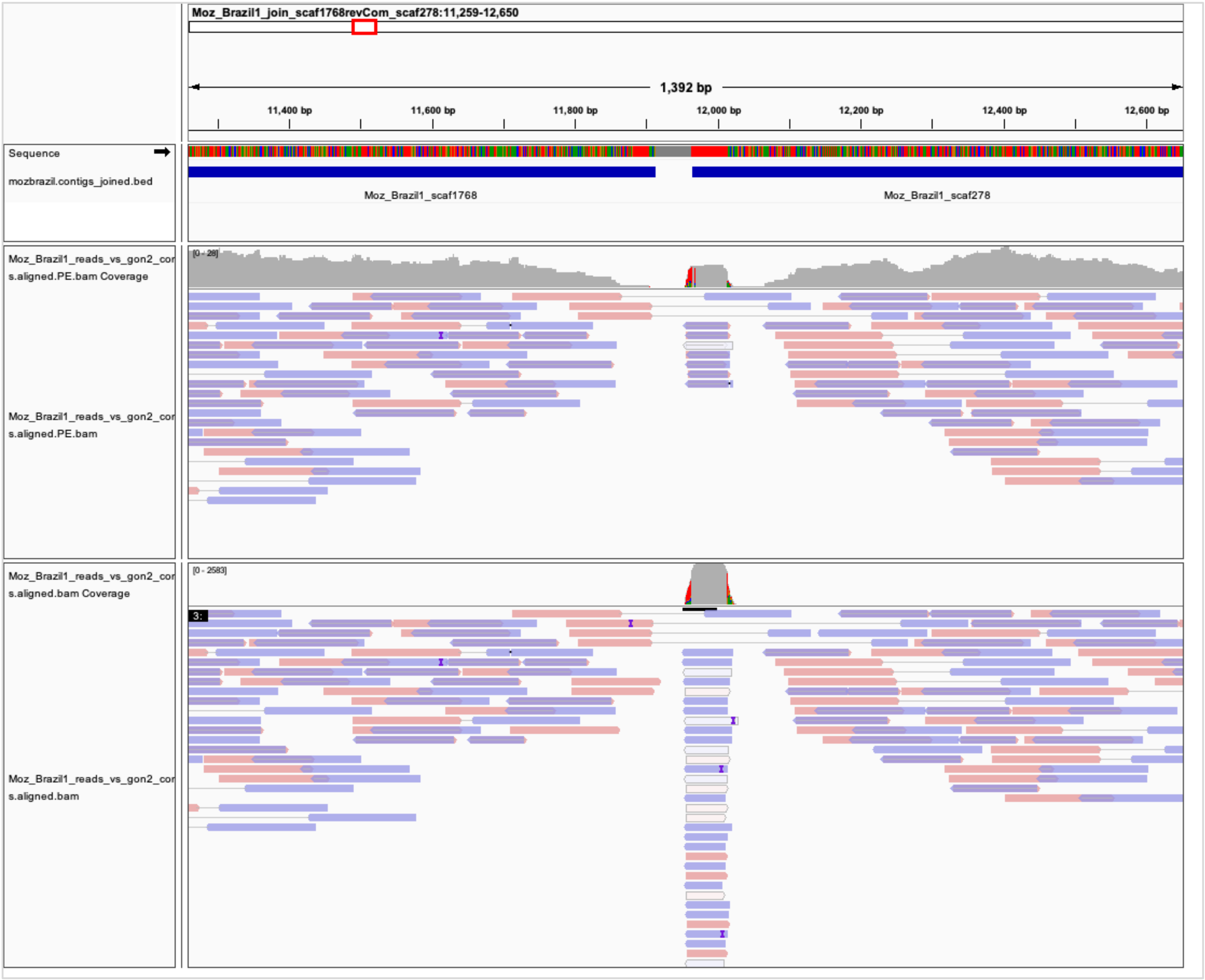

#### 1.8 Supplementary Figure 8. Read coverage analysis of the two contigs carrying the *gon-2* gene fragments in *M. ozzardi* isolate Moz-Venz-1.

The two contigs were tentatively joined in the orientation that would reconstruct an intact *gon-2* gene. The two contigs were separated by a stretch of 50 Ns, and Illumina reads used for assembly were mapped to the synthetic contig. The bottom track shows all reads. The top track shows only the subset of reads mapped as pairs.

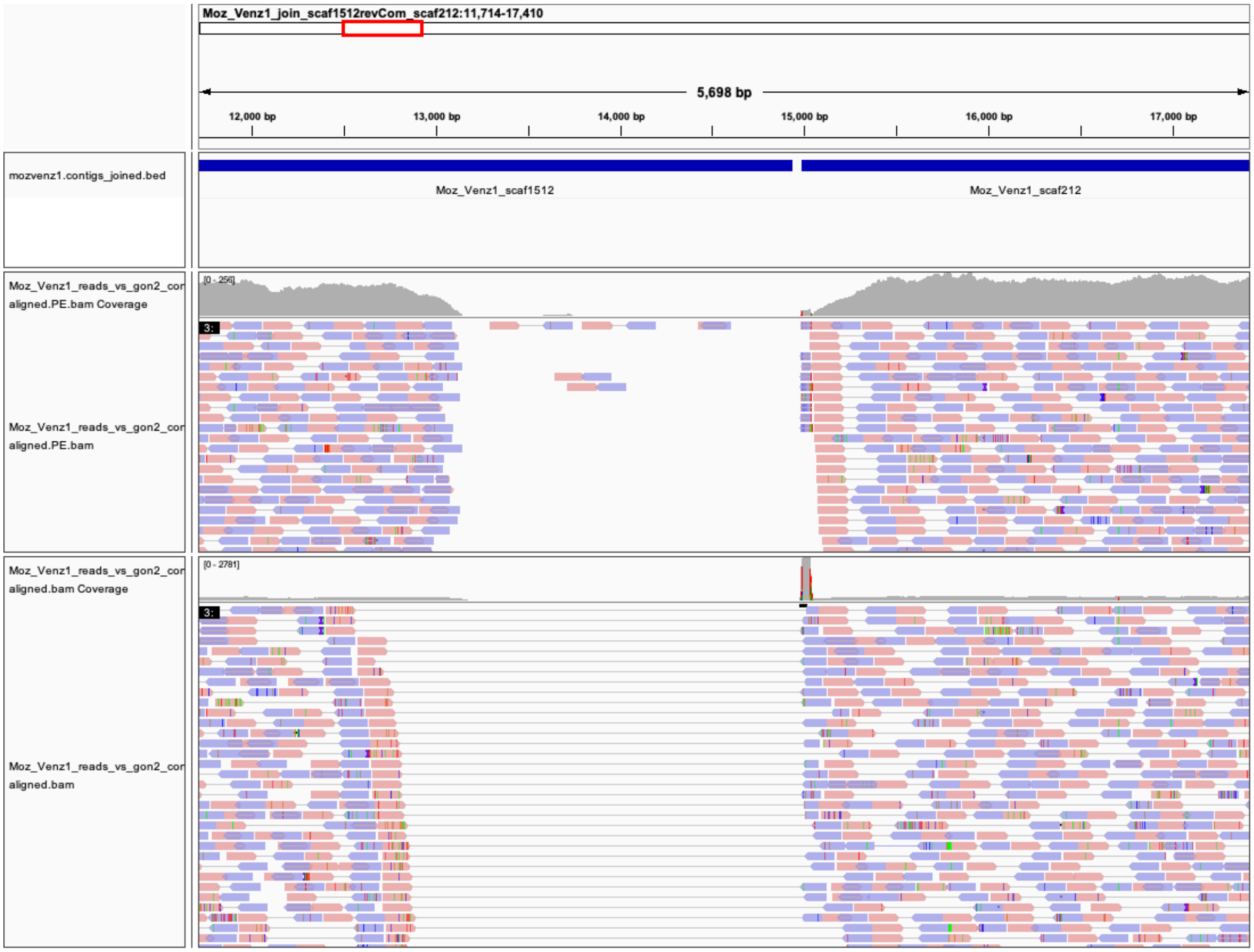

#### 1.9 Supplementary Figure 9. Read coverage analysis of the intact *gon-2* gene in *M. perstans* isolate Mpe-Cam-2.

Illumina reads used for assembly were mapped to the Mpe-Cam-2 contig carrying the intact *gon-2* gene. The region missing in *M. ozzardi gon-2* gene along with the flanking exons is marked in the red track. The bottom track shows all reads. The top track shows only the subset of reads mapped as pairs.

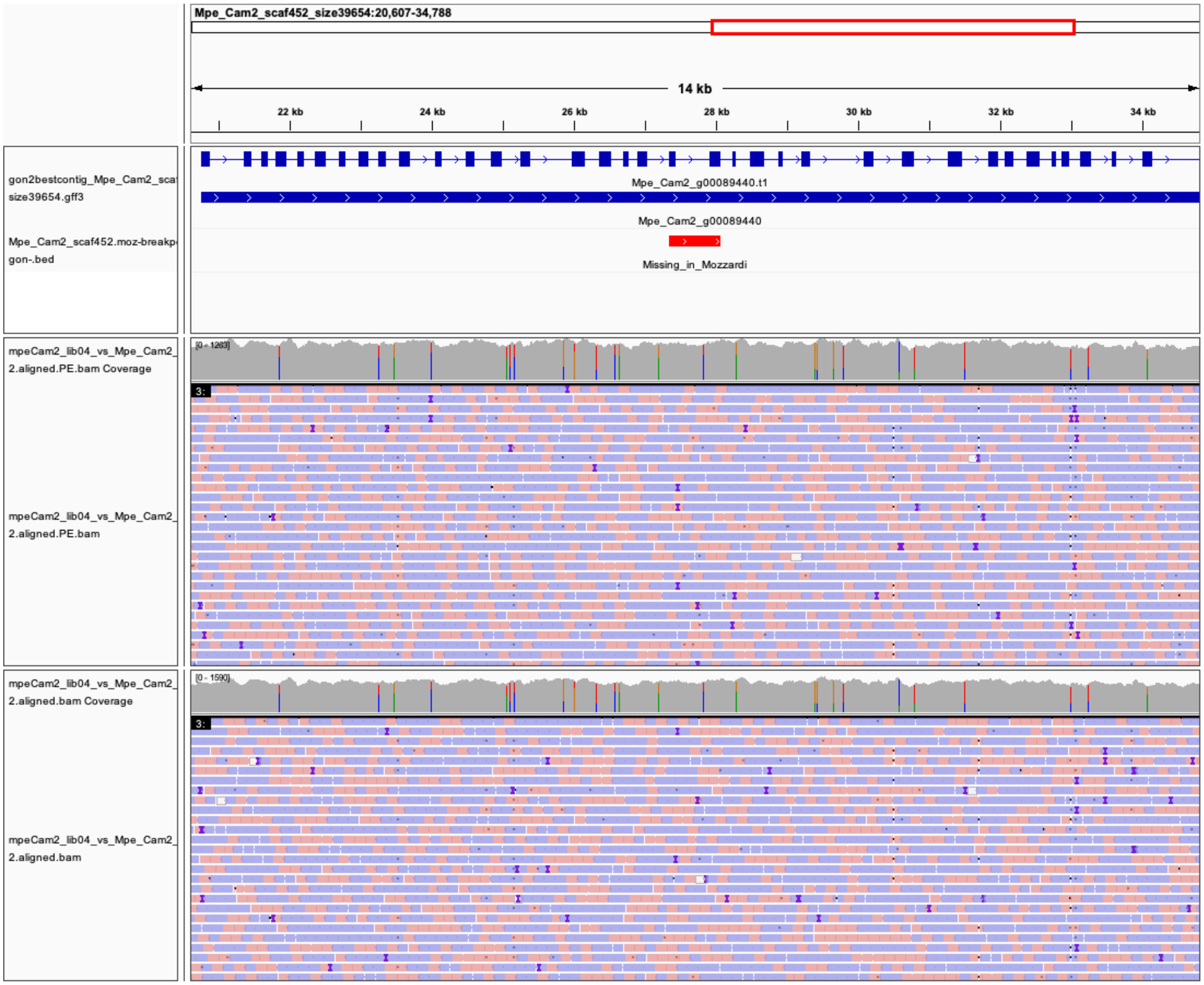

#### 1.10 Supplementary Figure 10. Phylogenetic analysis of DEC targets encoded by the *Cel-trp-1* , *Cel-trp-2* genes and their orthologs

Phylogenetic analysis of the *trp-* gene family was performed based on their protein sequences and the tree was rooted at the midpoint. Genes from *M. perstans* and *M. ozzardi* are in red, and *C. elegans* genes are in blue.

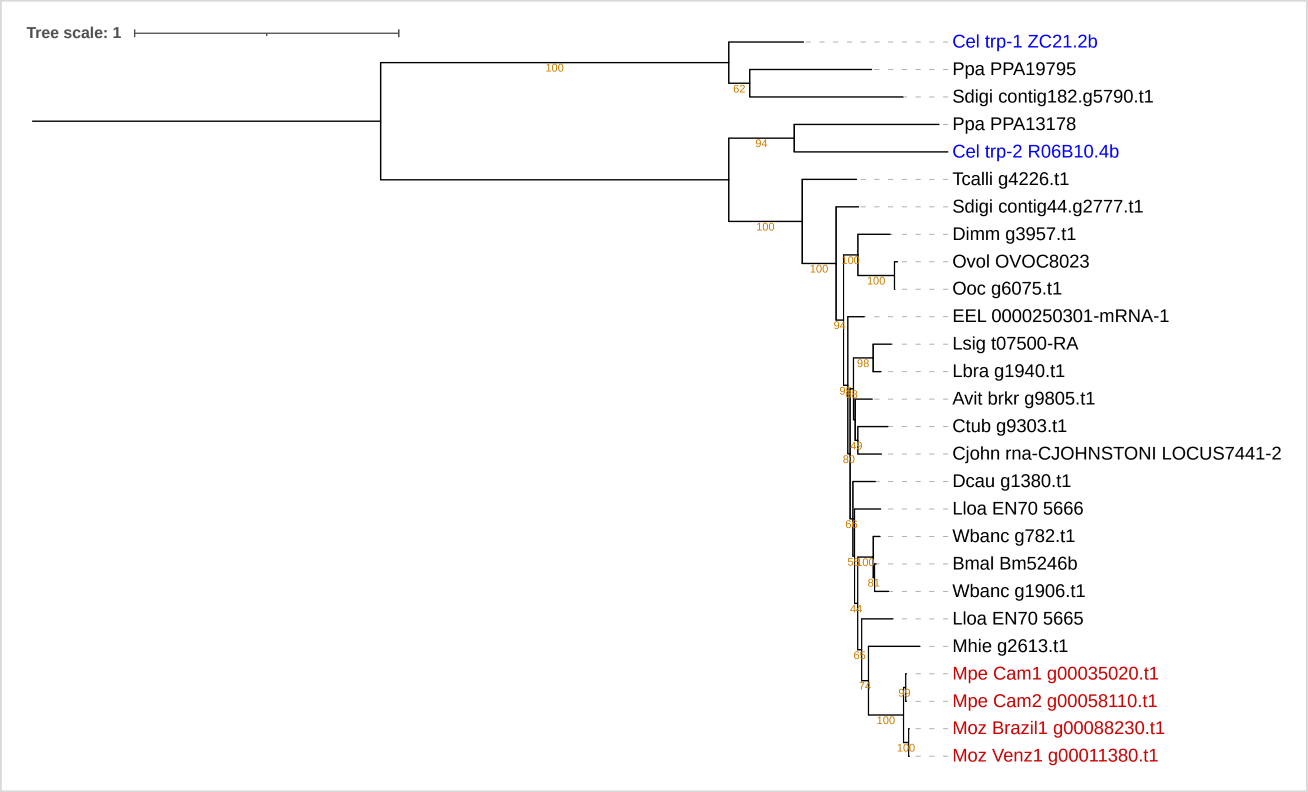

#### 1.11 Supplementary Figure 11. Phylogenetic analysis of DEC targets encoded by the *ced-11* gene and its orthologs

Phylogenetic analysis of the *ced-11* gene family was performed based on their protein sequences and the tree was rooted at the midpoint. Genes from *M. perstans* and *M. ozzardi* are in red, and *C. elegans* genes are in blue.

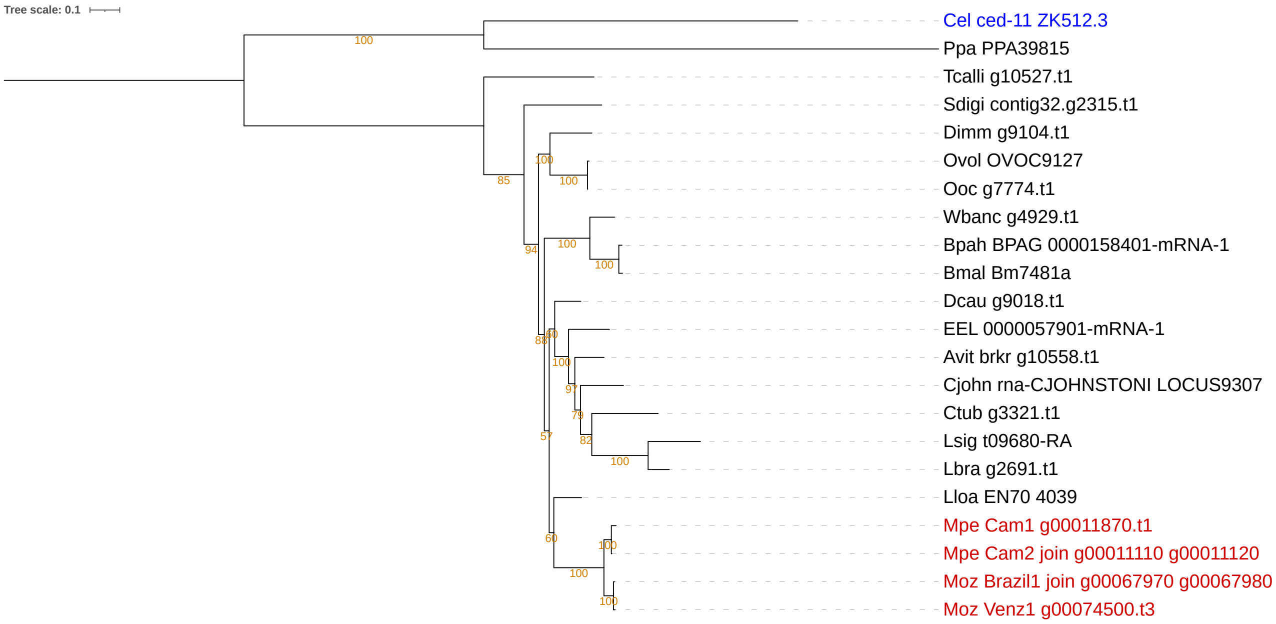

#### 1.12 Supplementary Figure 12. Phylogenetic analysis of DEC targets encoded by the *Cel-slo-1* gene and its orthologs

Phylogenetic analysis of the *slo-1* gene and its orthologs was performed based on their protein sequences and the tree was rooted at the midpoint. Genes from *M. perstans* and *M. ozzardi* are in red, and *C. elegans* genes are in blue.

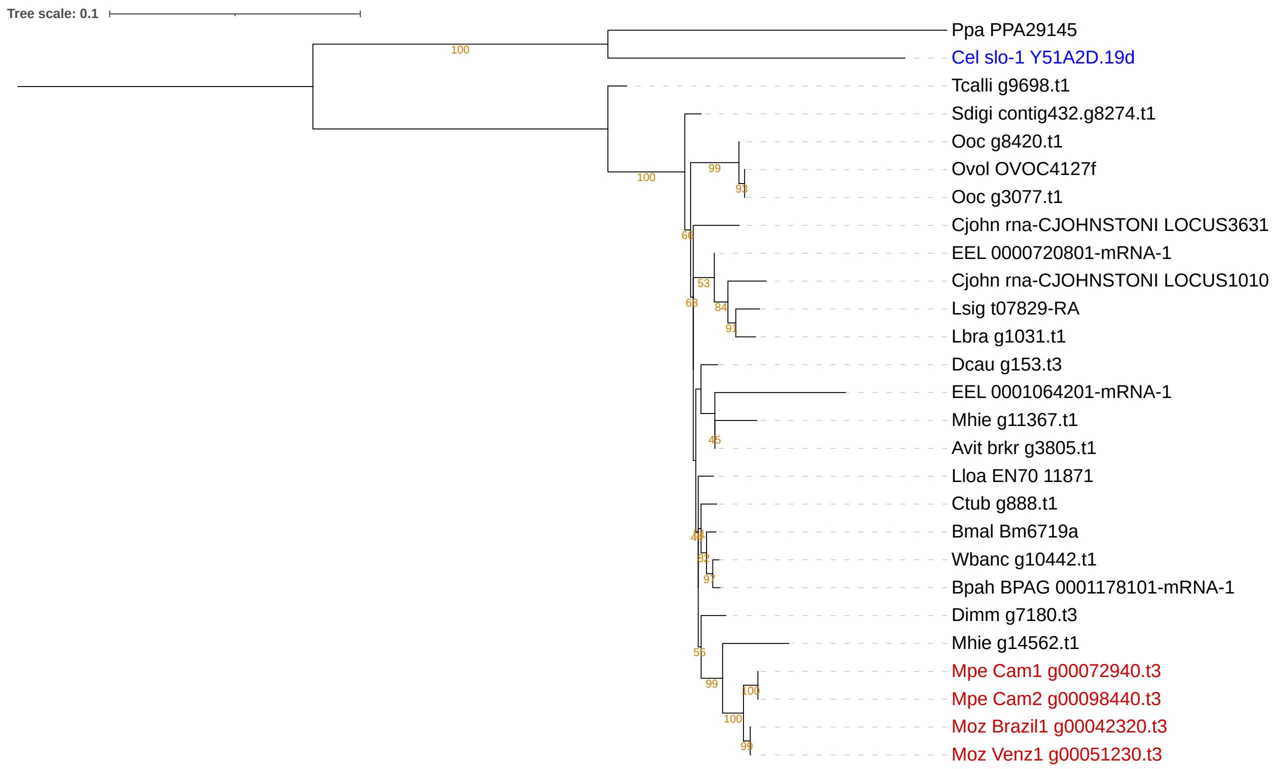

#### 1.13 Supplementary Figure 13. Phylogenetic analysis of *glc-4* gene and its orthologs

Phylogenetic analysis of the *glc-4* gene and its filarial orthologs was performed based on their protein sequences and the tree was rooted at the midpoint. Genes from *M. perstans* and *M. ozzardi* are in red, and *C. elegans* genes are in blue.

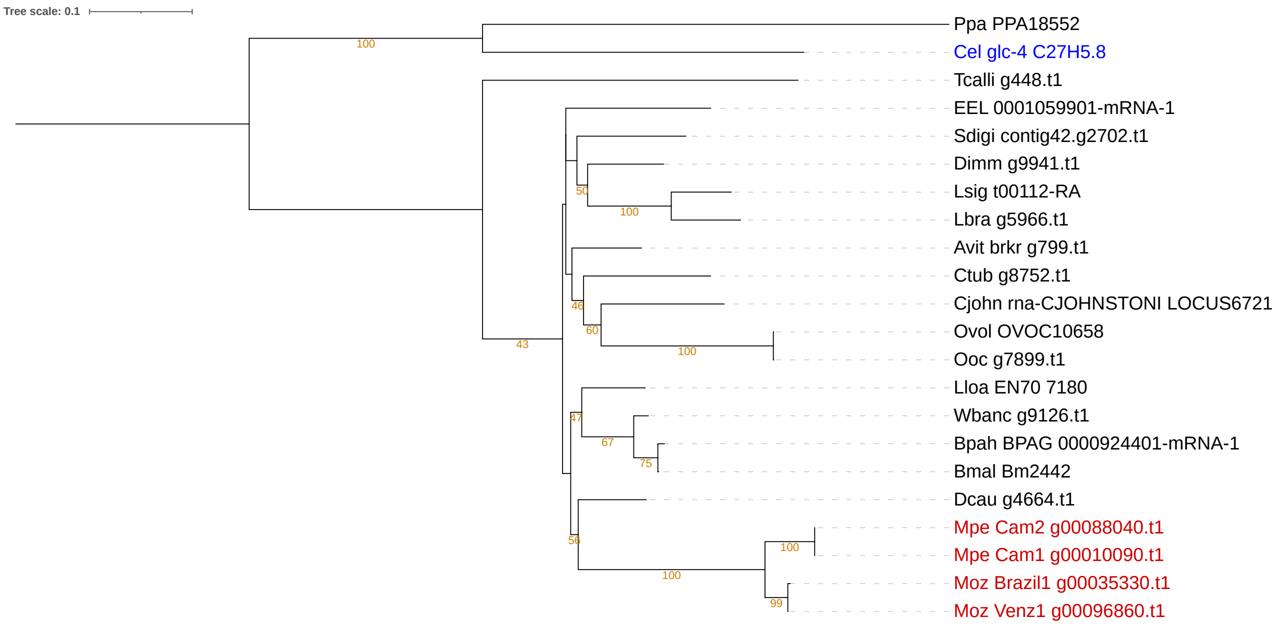

#### 1.14 Supplementary Figure 14. Phylogenetic analysis of the beta-tubulin gene family, that encodes various targets of albendazole

Phylogenetic analysis of the beta-tubulin gene family was performed based on their protein sequences and the tree was rooted at the midpoint. Genes from *M. perstans* and *M. ozzardi* are in red and *C. elegans* genes are in blue. Distinct clades that contain gene members from *Mansonella* species are highlighted in different background colors.

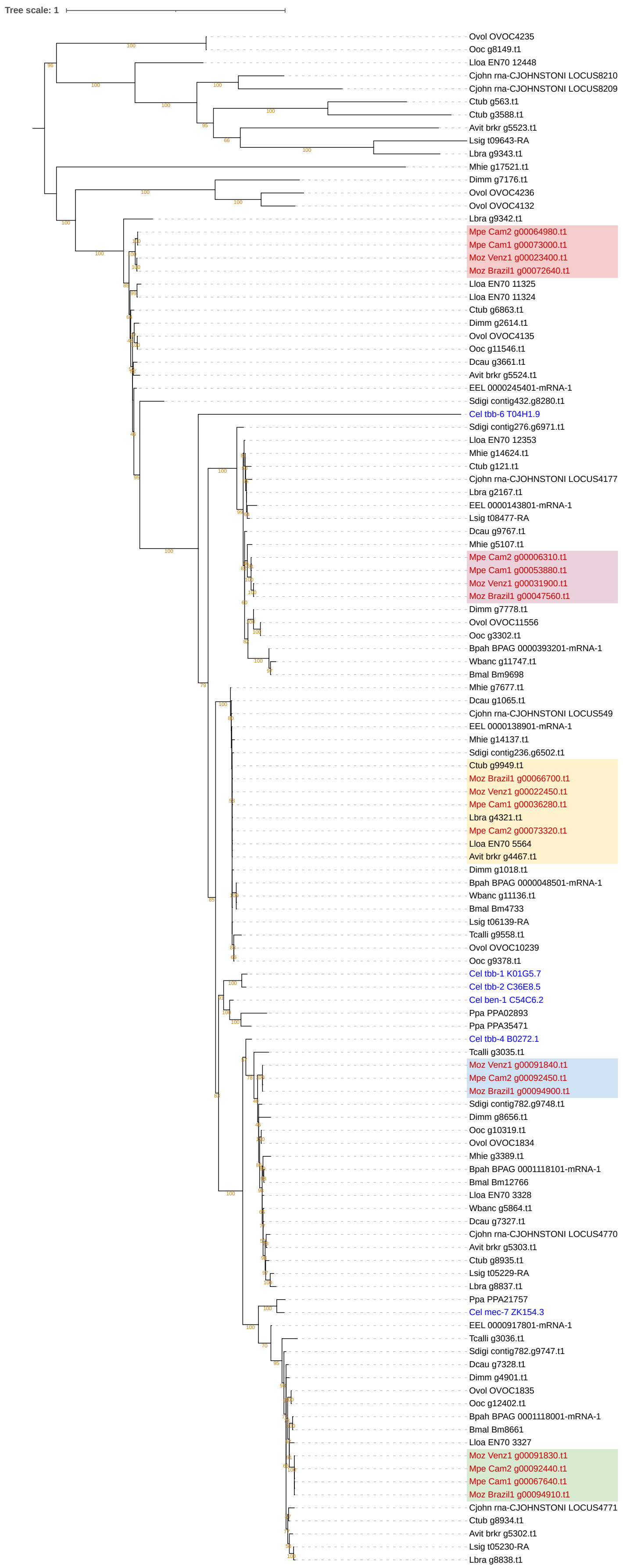
Tree scale: 1
Ovol OVOC4235
Ooc g8149.t1
Lloa EN70 12448
Cjohn rna-CJOHNSTONI LOCUS8210
Cjohn rna-CJOHNSTONI LOCUS8209
Ctub g563.t1
Ctub g3588.t1
Avit brkr g5523.t1
Lsig t09643-RA
Lbra g9343.t1
Mhie g17521.t1
Dimm g7176.t1
Ovol OVOC4236
Ovol OVOC4132
Lbra g9342.t1
Mpe Cam2 g00064980.t1
Mpe Cam1 g00073000.t1
Moz Venz1 g00023400.t1
Moz Brazil1 g00072640.t1
Lloa EN70 11325
Lloa EN70 11324
Ctub g6863.t1
Dimm g2614.t1
Ovol OVOC4135
Ooc g11546.t1
Dcau g3661.t1
Avit brkr g5524.t1
EEL 0000245401-mRNA-1
Sdigi contig432.g8280.t1
Cel tbb-6 T04H1.9
Sdigi contig276.g6971.t1
Lloa EN70 12353
Mhie g14624.t1
Ctub g121.t1
Cjohn rna-CJOHNSTONI LOCUS4177
Lbra g2167.t1
EEL 0000143801-mRNA-1
Lsig t08477-RA
Dcau g9767.t1
Mhie g5107.t1
Mpe Cam2 g00006310.t1
Mpe Cam1 g00053880.t1
Moz Venz1 g00031900.t1
Moz Brazil1 g00047560.t1
Dimm g7778.t1
Ovol OVOC11556
Ooc g3302.t1
Bpah BPAG 0000393201-mRNA-1
Wbanc g11747.t1
Bmal Bm9698
Mhie g7677.t1
Dcau g1065.t1
Cjohn rna-CJOHNSTONI LOCUS549
EEL 0000138901-mRNA-1
Mhie g14137.t1
Sdigi contig236.g6502.t1
Ctub g9949.t1
Moz Brazil1 g00066700.t1
Moz Venz1 g00022450.t1
Mpe Cam1 g00036280.t1
Lbra g4321.t1
Mpe Cam2 g00073320.t1
Lloa EN70 5564
Avit brkr g4467.t1
Dimm g1018.t1
Bpah BPAG 0000048501-mRNA-1
Wbanc g11136.t1
Bmal Bm4733
Lsig t06139-RA
Tcalli g9558.t1
Ovol OVOC10239
Ooc g9378.t1
Cel tbb-1 K01G5.7
Cel tbb-2 C36E8.5
Cel ben-1 C54C6.2
Ppa PPA02893
Ppa PPA35471
Cel tbb-4 B0272.1
Tcalli g3035.t1
Moz Venz1 g00091840.t1
Mpe Cam2 g00092450.t1
Moz Brazil1 g00094900.t1
Sdigi contig782.g9748.t1
Dimm g8656.t1
Ooc g10319.t1
Ovol OVOC1834
Mhie g3389.t1
Bpah BPAG 0001118101-mRNA-1
Bmal Bm12766
Lloa EN70 3328
Wbanc g5864.t1
Dcau g7327.t1
Cjohn rna-CJOHNSTONI LOCUS4770
Avit brkr g5303.t1
Ctub g8935.t1
Lsig t05229-RA
Lbra g8837.t1
Ppa PPA21757
Cel mec-7 ZK154.3
EEL 0000917801-mRNA-1
Tcalli g3036.t1
Sdigi contig782.g9747.t1
Dcau g7328.t1
Dimm g4901.t1
Ovol OVOC1835
Ooc g12402.t1
Bpah BPAG 0001118001-mRNA-1
Bmal Bm8661
Lloa EN70 3327
Moz Venz1 g00091830.t1
Mpe Cam2 g00092440.t1
Mpe Cam1 g00067640.t1
Moz Brazil1 g00094910.t1
Cjohn rna-CJOHNSTONI LOCUS4771
Ctub g8934.t1
Avit brkr g5302.t1
Lsig t05230-RA
Lbra g8838.t1
